# FlashS reveals multiscale spatial gene programs at atlas scale

**DOI:** 10.64898/2026.03.12.711372

**Authors:** Chen Yang, Xianyang Zhang, Jun Chen

## Abstract

Spatial transcriptomics links gene expression to tissue architecture, but detecting spatially variable genes at atlas scale remains difficult because biologically relevant patterns are multiscale, sparse, and often non-parametric. Here we show that FlashS, a frequency-domain kernel test using random Fourier features, sparse sketching, and a kurtosis-corrected null, retains Gaussian-kernel flexibility at near-linear per-gene cost without permutation. Across 50 benchmark datasets spanning 9 spatial transcriptomics platforms, FlashS achieves the highest ranking accuracy among 14 compared methods while maintaining calibrated inference under simulation and full-atlas permutation. In human heart tissue, it recovers a mitochondrial biogenesis program that co-localizes with ventricular cardiomyocytes. On the Allen Brain MERFISH atlas containing 3.94 million cells, FlashS completes in minutes on a single workstation and separates biological signal from negative-control barcodes even when p-values saturate.

## Introduction

Spatial transcriptomics technologies now measure gene expression in its tissue context, turning spatially variable gene (SVG) detection into a first step for discovering how tissue architecture organizes biological function. SVGs encode morphogen gradients in development, metabolic zonation, immune infiltration at tissue interfaces, and regionalized transcriptional programs in the brain. A useful SVG test must therefore do more than rank genes by smoothness: it must detect focal, gradient, periodic, boundary, and compound patterns while controlling false discoveries on sparse count data.

This requirement exposes a basic statistical and computational tension. Gaussian process (GP)-based methods[1–4] provide flexible spatial models, but their covariance matrices are quadratic or cubic in the number of locations. Scalable methods avoid this cost by narrowing the representation: covariance projection in SPARK-X[5], granularity tests in scBSP[6], local autocorrelation in Moran’s I and Hotspot[7], or fixed B-spline bases in PreTSA[8]. These restrictions are computationally effective, but they make the test less sensitive to patterns outside the chosen basis. The problem is amplified in sequencing-based spatial transcriptomics, where 80–95% zero entries per gene are common and the zero component itself can carry spatial information.

FlashS starts from a simple premise: if spatial expression is an unknown function over tissue coordinates, the computational bottleneck should be removed from the kernel rather than from the class of patterns the kernel can represent. Random Fourier Features (RFF)[9] approximate Gaussian kernels through inner products in a spectral feature space, eliminating the *n* × *n* matrix while preserving a broad, non-parametric spatial basis. This distinguishes FlashS from two related strategies. Graph-based spectral methods such as SpaGFT[10] decompose expression onto eigenvectors of a spatial graph Laplacian; because the spectrum is discrete and graph-dependent, the available frequencies are fixed by a particular neighbourhood construction. Low-rank or inducing-point GP approximations[3] replace the full kernel with a coarsened grid, smoothing fine-scale patterns below the grid spacing. RFF instead samples frequencies from the *continuous* spectral density of the Gaussian kernel, so local foci, tissue-wide gradients, and multi-scale mixtures all contribute evidence without requiring a specific graph or grid. Because each gene is projected through its non-zero entries, sparsity reduces computation instead of forcing dense preprocessing.

The resulting framework combines three elements designed for the data-generating features of spatial transcriptomics. Adaptive bandwidths span local neighbourhoods to tissue-wide scales using the median nearest-neighbour distance and the Fiedler value[11] of the spatial graph. A three-channel test separates presence–absence, rank-transformed intensity, and raw-count signal, then combines evidence across channels and scales by the Cauchy rule[12]. Finally, a kurtosis-corrected Satterthwaite null accounts for correlated RFF columns and heavy-tailed zero-inflated expression, giving analytic p-values without permutation.

We evaluate FlashS (**F**requency-domain **L**arge-scale **A**nalysis of **S**patial **H**eterogeneity) as both a statistical test and a biological discovery tool. Across 50 benchmark datasets spanning 9 spatial transcriptomics platforms[13], FlashS achieves the highest SVG ranking accuracy. Its additional detections are not merely more numerous: in brain and heart tissue they are enriched for interpretable spatial programs, including a PGC-1*α*-regulated mitochondrial biogenesis program in human heart that co-localizes with ventricular cardiomyocytes. At the other end of the scale, FlashS analyses the 3.94-million-cell Allen Brain MERFISH atlas in minutes on standard hardware with calibrated inference under full-atlas permutation.

## Results

### Frequency-domain testing for sparse multi-scale spatial expression

For each gene, FlashS converts spatial coordinates into a *D*-dimensional random Fourier feature matrix **Z** whose inner products approximate a Gaussian kernel, and tests the total squared projection *T* =∥ **Z**^⊤^**y** ∥^2^ of expression onto these features (Fig. 1). Sparse sketching and implicit centering keep the per-gene cost at *O*(nnz · *D*) with no *n* × *n* matrix materialized. Three complementary channels—binary, rank-transformed, and raw counts—are evaluated at each of *L* bandwidth scales, yielding 3*L* + 3 p-values (including linear trend tests) that are combined by the Cauchy rule into a single gene-level p-value against the kurtosis-corrected moment-matching null.

**Fig. 1.**
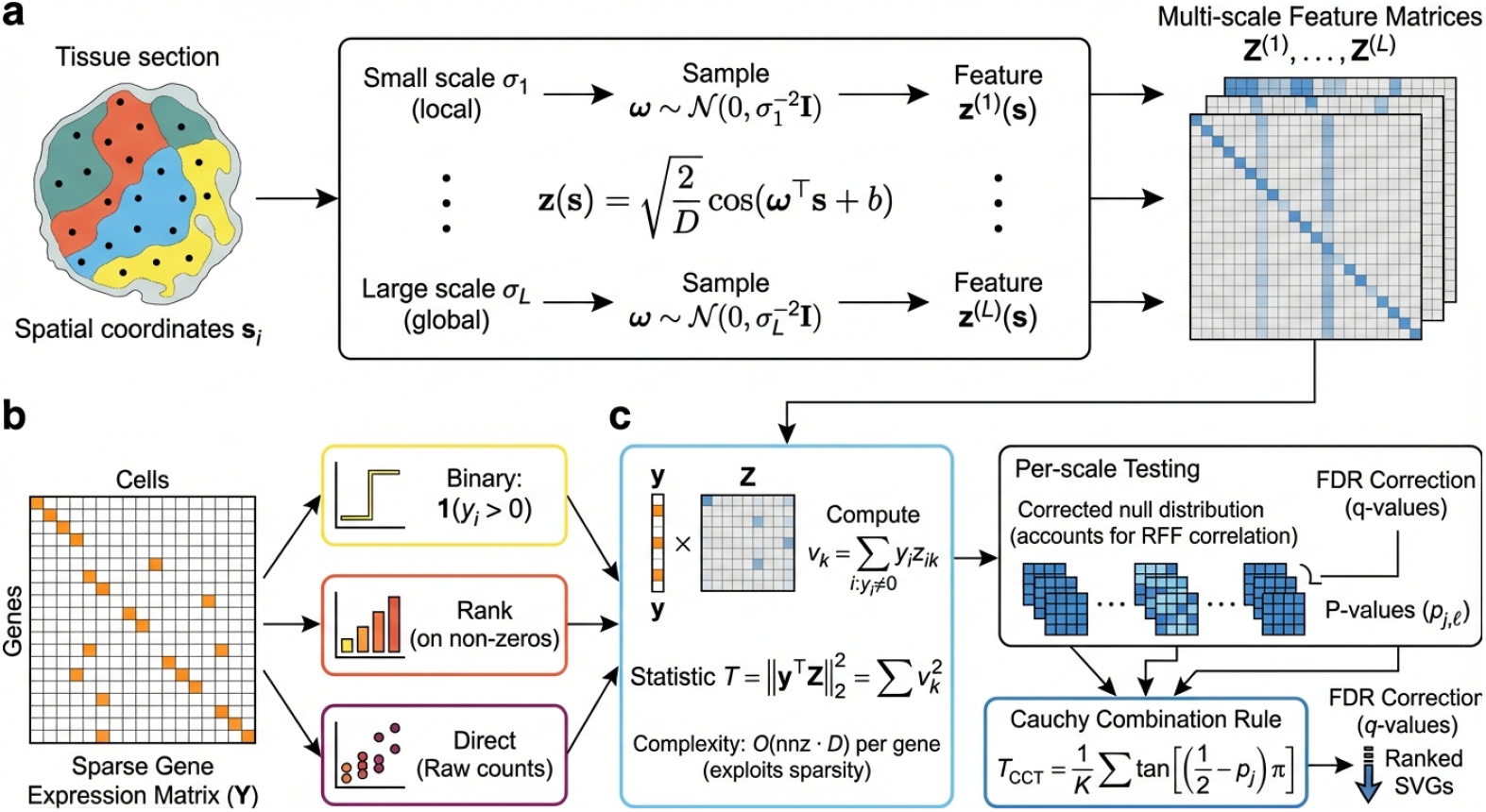
Overview of. FlashS. **a**, Spatial coordinates are transformed into Random Fourier Features (RFF), approximating kernel evaluations via inner products in a low-dimensional feature space. **b**, A three-part test evaluates binary expression, rank-transformed intensities, and raw counts against spatial features at multiple bandwidth scales. **c**, The test statistic *T* = ∥ **Z**^⊤^**y** ∥^2^ aggregates squared projections of expression onto spatial features. Per-scale p-values are combined via the Cauchy combination rule to yield a single p-value per gene.

The framework has two tuning parameters, the RFF feature count *D* and the number of bandwidth scales *L* (defaults *D* = 500, *L* = 7); because *D* is fixed, changing *L* redistributes features across scales without affecting per-gene cost. FlashS operates on raw counts, which preserves the zero component that library-size normalization eliminates; ablations supporting this choice and sensitivity to *D* and *L* are reported in Supplementary Fig. 1.

### Benchmark accuracy on the Open Problems SVG evaluation

We evaluated FlashS on the Open Problems benchmark for SVG detection[13], a standardized evaluation framework comprising 50 datasets across 9 spatial transcriptomics platforms (Visium, Slide-seq V2, MERFISH, STARmap, Stereo-seq, seqFISH, DBiT-seq, Slide-tags, and Post-Xenium). Datasets span diverse organisms (human, mouse, *Drosophila*) and tissues (brain, cancer, embryo, heart, kidney, and others), with sizes ranging from 936 to 29,309 spots and 210 to 1,050 genes per dataset. Ground truth spatial variability scores were generated using scDesign3[14] by mixing Gaussian process and non-spatial models. Performance is measured by Kendall *τ* correlation between predicted and true spatial variability scores, averaged across genes grouped by their original feature identity.

FlashS achieves a mean Kendall *τ* of 0.935 across all 50 datasets, the highest accuracy among the compared methods (Fig. 2; Supplementary Table 1). This surpasses the previous best method, SPARK-X (*τ* = 0.886), by Δ*τ* = 0.049. We independently validated this comparison by running SPARK-X in the same computing environment on all 50 datasets, obtaining *τ* = 0.881 (consistent with the leaderboard value), with FlashS outperforming on every dataset (Δ*τ* = +0.054; Supplementary Table 10). Notably, FlashS maintains positive correlation across all 50 datasets (minimum *τ* = 0.788), whereas several competing methods produce negative correlations on individual datasets, indicating anti-correlated predictions. Performance is consistent across all 9 platforms, with mean *τ* > 0.80 on every platform (Supplementary Fig. 2).

**Fig. 2.**
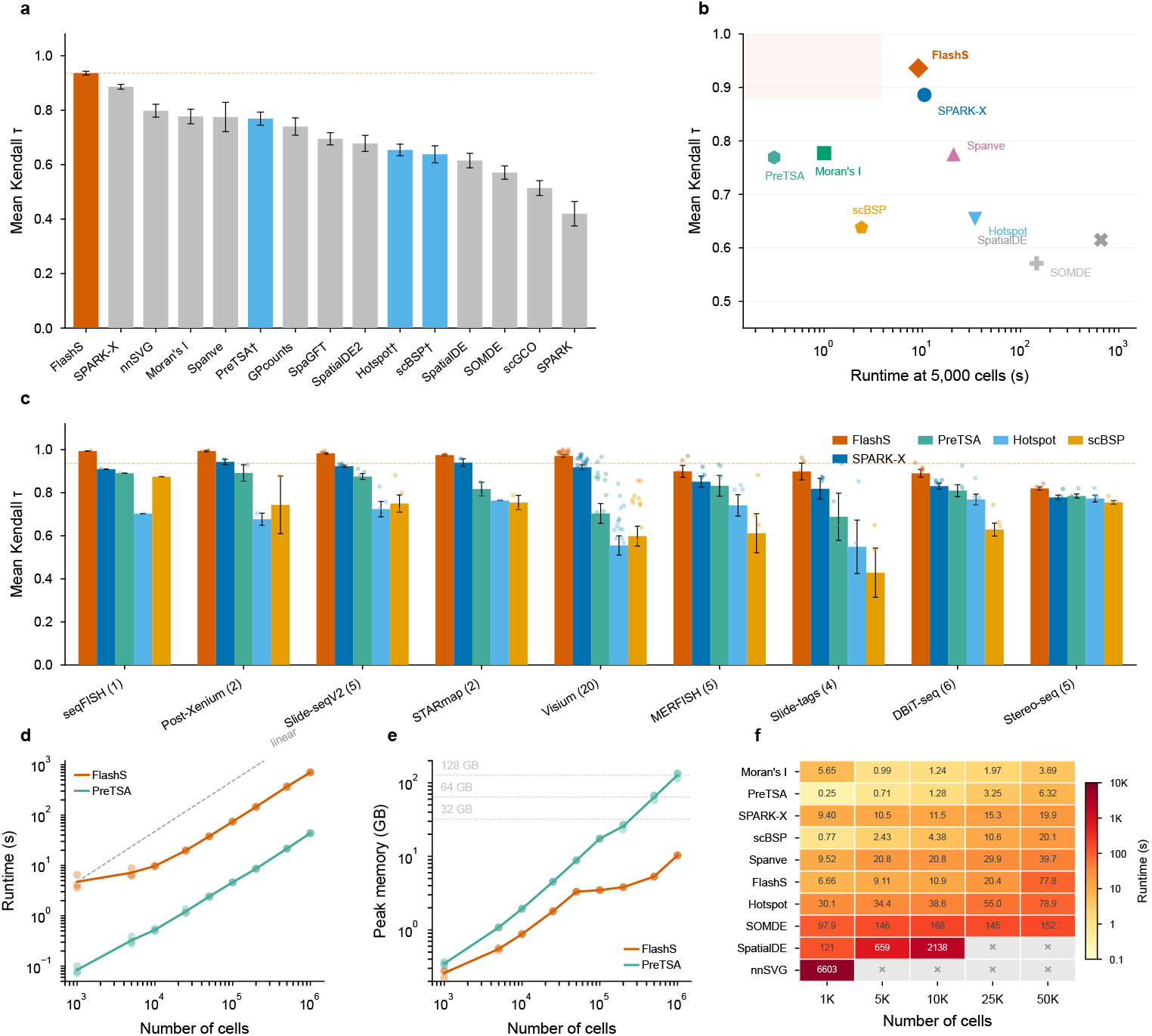
Benchmark accuracy, scalability, and per-platform performance. **a**, Mean Kendall *τ* correlation between predicted and ground-truth spatial variability scores across 50 datasets spanning 9 platforms. FlashS (red) achieves *τ* = 0.935, surpassing SPARK-X (*τ* = 0.886) by Δ*τ* = 0.049. PreTSA, Hotspot and scBSP (marked with *†*) were benchmarked independently on the same datasets. Error bars indicate standard error of the mean across biologically independent benchmark datasets. **b**, Accuracy versus runtime (at 5,000 cells). FlashS occupies the Pareto-optimal region of high accuracy and fast runtime (shaded). **c**, Per-platform mean Kendall *τ* for FlashS, SPARK-X, PreTSA, Hotspot, and scBSP across 9 platforms, sorted by FlashS performance. Individual dataset values are shown as dots; error bars indicate standard error of the mean. The numbers of biologically independent datasets are *n* = 20 (Visium), *n* = 6 (DBiT-seq), *n* = 5 each (MERFISH, Slide-seq V2, and Stereo-seq), *n* = 4 (Slide-tags), *n* = 2 each (Post-Xenium and STARmap), and *n* = 1 (seqFISH). **d**, Runtime as a function of cell number (5,000 genes). Both FlashS and PreTSA scale near-linearly; dashed line indicates linear scaling. **e**, Peak memory usage. PreTSA’s memory scales as *O*(*n · g*), crossing 32 GB at *∼*200,000 cells and reaching 137 GB at 1,000,000 cells; FlashS remains below 11 GB via sparse sketching. Dotted lines mark typical workstation memory. Points show individual independent computational replicates (*n* = 3 per size). **f**, Runtime heatmap for 10 SVG detection methods across cell counts (1,000–50,000), sorted by speed at 50,000 cells. Cell values show mean runtime in seconds across *n* = 3 independent computational replicates; *×* marks timeout (>2 h) or out-of-memory failure. All benchmarks were performed on a 16-core compute node.

To contextualize the accuracy–scalability trade-off, we additionally benchmarked three scalable methods not included in the original Open Problems leaderboard— PreTSA[8], Hotspot[7], and scBSP[6]—on the same 50 datasets (Fig. 2a). PreTSA, which applies B-spline tensor product regression with an F-test, achieves mean *τ* = 0.769; Hotspot achieves mean *τ* = 0.654 and scBSP mean *τ* = 0.638. Despite its computational efficiency, PreTSA’s parametric regression captures smooth spatial trends but cannot model the multi-scale, non-smooth patterns that kernel-based methods detect, resulting in a Δ*τ* = 0.166 gap relative to FlashS. On the accuracy–runtime Pareto frontier (Fig. 2b), FlashS is more accurate than other scalable alternatives and faster than methods of comparable accuracy.

### Scalability to million-location datasets

Emerging spatial atlases profile millions of cells across entire organs, demanding SVG detection methods that scale without sacrificing accuracy. We benchmarked FlashS on simulated datasets with 5,000 genes and cell counts ranging from 1,000 to 1,000,000 (Fig. 2d,e). Runtime scales near-linearly with cell number, reaching 11.7 minutes at one million cells with peak memory of 10.3 GB—within the capacity of a standard workstation (Supplementary Table 2).

This memory efficiency distinguishes FlashS from the only other method of comparable accuracy. Among ten compared methods (Fig. 2f; Supplementary Table 3), PreTSA achieves the lowest wall-clock time but its memory scales as *O*(*n* · *g*), reaching 137 GB at one million cells—13-fold more than FlashS (Supplementary Table 4). On a typical workstation (32–64 GB RAM), PreTSA becomes infeasible beyond ∼200,000 cells. GP-based methods (SpatialDE, nnSVG) exceed the 2-hour timeout at ≥5,000 cells (Supplementary Table 3). Among methods with competitive accuracy (*τ* > 0.85), FlashS is the fastest.

### Statistical validity and detection profile

Accurate inference under the null is the foundation for any SVG test: p-values must be calibrated for false discoveries to be controlled, and the test must retain power across the spatial pattern types that real tissue contains. Two features of spatial transcriptomics complicate this task—the RFF features are correlated, particularly at large bandwidth scales, and zero-inflated expression vectors are heavy-tailed, violating the Gaussian assumption of standard quadratic-form approximations. We derive a kurtosis-corrected moment-matching null that accounts for both effects (Methods; Supplementary Note 3). Ablation analysis confirms that both corrections are required: the naive *χ*^2^ baseline that ignores RFF covariance inflates the FPR to 33% at *α* = 0.05 (6.6 × nominal), Gaussian covariance correction alone brings it to 6.6% at 95% sparsity, and the kurtosis term reduces it further to 4.4%—with the gap widening in the extreme-tail regime (*α* = 10^−3^: naive 25%, cov-only 0.28%, kurtosis-corrected 0.17%; Supplementary Fig. 3a). To assess calibration we simulated null datasets on 2,500-spot grids at three zero-fraction levels (∼50%, ∼90%, ∼95%; 20 replicates × 1,000 genes). With the corrected null, the empirical FPR at *α* = 0.05 averages 5.6% across 60 replicates (Fig. 3a; Supplementary Fig. 4b), close to the nominal level; the q-value-based false discovery rate[15] remained below 0.1% at *q* < 0.05 across all 60 replicates, and per-scale calibration is accurate at every bandwidth, including the largest scales where RFF columns are most correlated (Supplementary Fig. 11a).

**Fig. 3.**
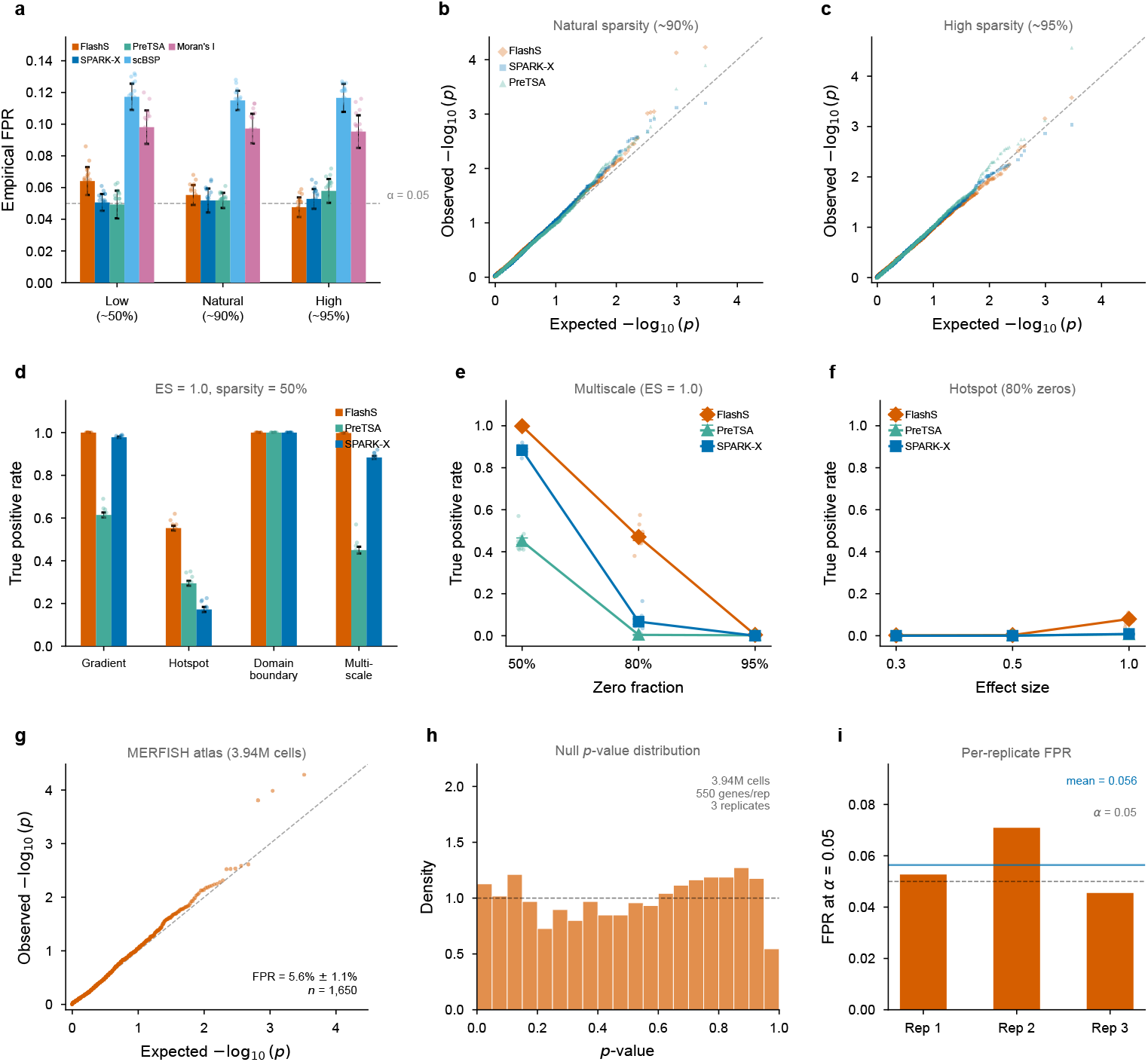
Statistical calibration and detection power. **a**, Empirical false positive rates at *α* = 0.05 across five methods and three sparsity levels (*∼*50%, *∼*90%, *∼*95% zeros). Bars show the mean, error bars show standard deviation, and dots show *n* = 20 independent simulation replicates of 1,000 genes each. The dashed line marks the nominal level. FlashS, SPARK-X, and PreTSA track the nominal level; scBSP and Moran’s I show inflated FPR. **b,c**, QQ plots of null p-values at natural (*∼*90%) and high (*∼*95%) sparsity from the same simulations. FlashS, SPARK-X, and PreTSA track the diagonal closely. **d**, Cross-method detection power (TPR at Benjamini–Hochberg *q* < 0.05) across four spatial pattern types at effect size 1.0 and 50% sparsity. FlashS achieves the highest sensitivity on hotspot and multiscale patterns. **e**, TPR as a function of sparsity for the multiscale pattern (effect size 1.0). At 80% zeros, FlashS retains substantial power (47%) while SPARK-X and PreTSA drop to 7% and <1%. **f**, TPR as a function of effect size for the hotspot pattern at 80% sparsity. In **d–f**, lines or bars show the mean, error bars show the standard error of the mean, and translucent dots show *n* = 10 independent simulation replicates (200 spatial and 200 null genes per replicate). **g**, QQ plot of null p-values from global row permutation on the Allen MERFISH atlas (3.94 million cells, 550 genes, *n* = 3 independent permutations pooled, 1,650 null tests). FlashS maintains near-nominal FPR at million-cell scale (mean FPR = 5.6% *±* 1.1% s.d.). **h**, Histogram of MERFISH null p-values with uniform reference density. **i**, False positive rate for each of the *n* = 3 independent permutations at *α* = 0.05; the blue line is the arithmetic mean and the dashed line is the nominal level.

Cross-method comparison reveals a range of calibration quality (Fig. 3a). FlashS (4.8–6.4%), SPARK-X (5.1–5.3%), and PreTSA (4.9–5.8%) all track the nominal level closely, while scBSP (11.5–11.7%) and Moran’s I (9.5–9.8%) show inflated Type I error. Good calibration does not guarantee detection power across diverse spatial pattern types. To characterise detection profiles, we simulated four canonical spatial pattern types—hotspot (localized expression), gradient (tissue-wide trend), multiscale (compound), and domain (sharp boundaries)—at varying effect sizes and sparsity levels (Fig. 3d–f; Supplementary Fig. 4c). At effect size 1.0 and 50% sparsity, FlashS achieves the highest sensitivity on hotspot and multiscale patterns. On hotspot patterns, FlashS achieves TPR = 55.3% versus 29.4% for PreTSA and 17.2% for SPARK-X. On domain patterns, all three methods achieve 100% power; on gradient patterns, FlashS (100%) and SPARK-X (97.8%) detect nearly all SVGs while PreTSA reaches 61.4%. At higher sparsity (80% zeros), the advantage widens on multiscale patterns: FlashS retains TPR = 47.1% while SPARK-X drops to 6.7% and PreTSA to 0.4%. These detection gaps predict that single-basis methods will miss genes with multi-scale or localized spatial structure in real tissue—a prediction tested in the biological analyses below.

To confirm that calibration extends to atlas-scale data with realistic spatial geometry, we performed global row permutation testing on the full Allen MERFISH dataset (3.94 million cells, 550 genes). Global row permutation destroys all spatial structure while preserving the real coordinate layout and per-gene marginal distributions. Across three independent replicates (1,650 total null tests), FlashS maintained a mean FPR of 5.6% ± 1.1% at *α* = 0.05 (Fig. 3g–i), confirming calibrated inference at million-cell scale on real spatial coordinates. Complementary structured permutations that selectively preserve part of the spatial organization—within-section shuffling and spatial block permutation—yield 100% rejection across all replicates (Supplementary Table 9), so detections track the degree of preserved spatial organization rather than residual technical structure.

We next asked whether the additional sensitivity of FlashS translates into reproducible biological signal. For this analysis we emphasized two anchor baselines representing distinct modelling choices: SPARK-X, the strongest non-parametric benchmark competitor (*τ* = 0.886), and PreTSA, the leading parametric regression-based alternative (*τ* = 0.769). SPARK-X tests whether a richer spectral basis adds biological information beyond a calibrated kernel-projection detector; PreTSA tests whether non-parametric multi-scale patterns recover biology missed by a fixed smooth basis.

### Reproducibility across samples and platforms

To assess cross-sample reproducibility, we compared FlashS with SPARK-X on 12 Visium samples from human dorsolateral prefrontal cortex (3 donors, 33,538 genes; Maynard et al.[16]). FlashS rankings were highly concordant across biological replicates (same-donor mean *τ* = 0.703, cross-donor mean *τ* = 0.643; Fig. 4a), with inter-individual variation exceeding within-donor variation as expected. Both methods detected all 27 curated cortical layer markers at *q* < 0.05, and both enriched these markers among top-ranked genes (top-100: SPARK-X 33.7×, FlashS 29.0×; Fig. 4b,c; Supplementary Fig. 9). Thus, FlashS preserves recovery of canonical spatial biology rather than trading precision for a larger call set.

**Fig. 4.**
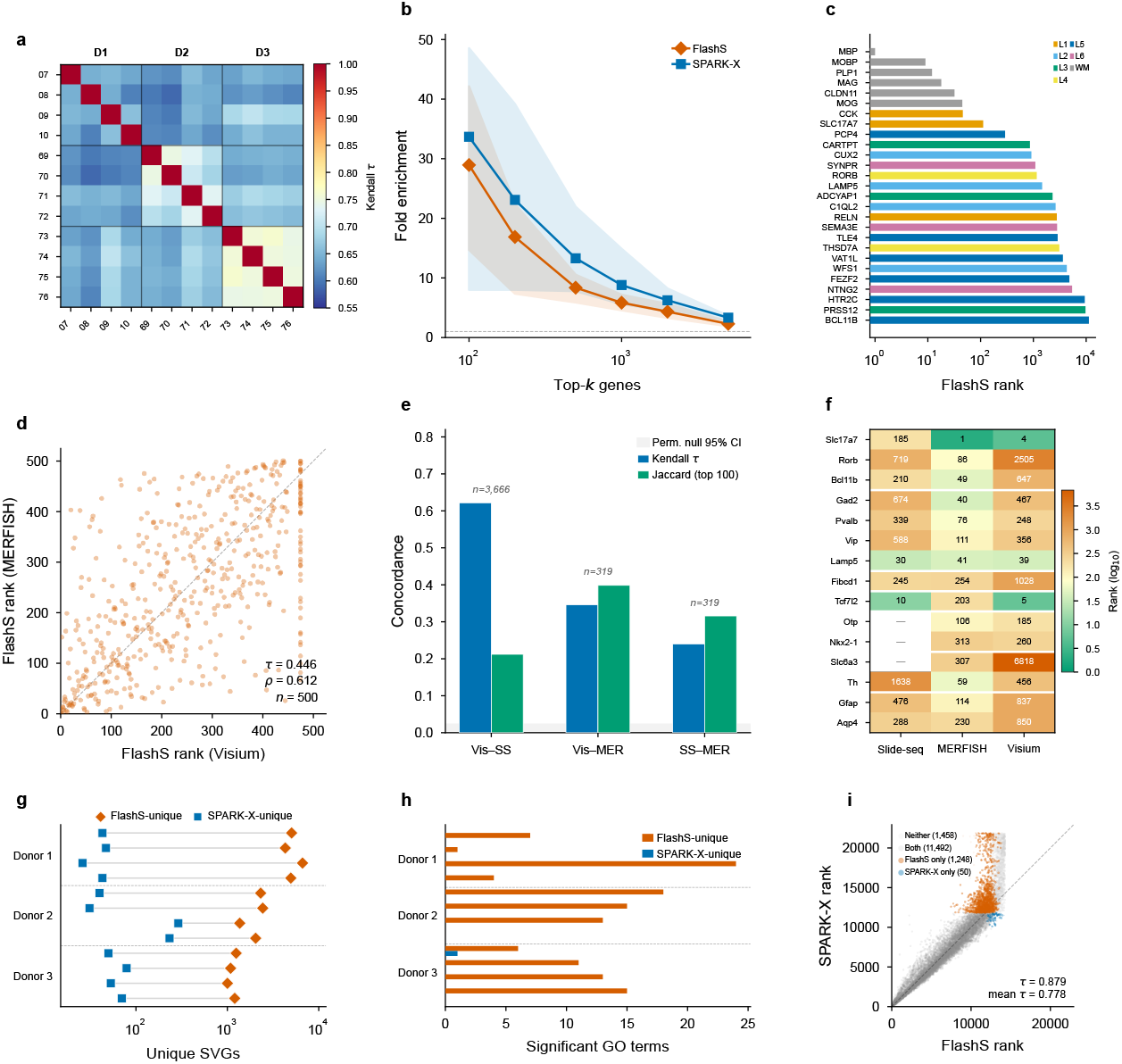
Cross-dataset and cross-platform generalization of SVG rankings. **a**, Cross-sample reproducibility across 12 biologically independent DLPFC Visium samples from 3 donors. Same-donor pairs show higher concordance (mean *τ* = 0.703) than cross-donor pairs (mean *τ* = 0.643). **b**, Fold enrichment of 27 cortical-layer markers in top-*k* genes across the same *n* = 12 samples. Lines are arithmetic means; bands are 95% confidence intervals of the mean. **c**, FlashS ranks of the 27 markers, colored by layer. All are SVGs (*q* < 0.05). **d**, Rank concordance between Visium mouse brain and MERFISH for 500 common genes (Kendall *τ* = 0.446, Spearman *ρ* = 0.612, two-sided permutation *P* < 0.001, 1,000 permutations). **e**, Pairwise Kendall *τ* and top-100 Jaccard similarity across Visium, Slide-seq V2, and MERFISH. The grey band spans the 2.5th–97.5th percentiles of the two-sided null from 1,000 permutations. Visium–Slide-seq uses 3,666 genes (*τ* = 0.622); Visium–MERFISH and Slide-seq–MERFISH use 319 genes (*τ* = 0.346 and 0.240). **f**, Ranks of 15 prespecified brain-region markers across platforms. **g**, Method-specific SVG counts across the *n* = 12 DLPFC samples after Benjamini–Hochberg correction at *q* < 0.05. Lines connect paired sections; mean unique counts are 2,802 for FlashS and 84 for SPARK-X. **h**, Significant Gene Ontology terms for method-specific SVGs across the same samples. FlashS-unique SVGs yield 0–24 terms; SPARK-X-unique SVGs yield zero in 11 of 12 samples. **i**, Rank comparison on sample 151673 (14,248 common genes); categories use Benjamini–Hochberg *q* < 0.05.

The difference appears in the genes beyond the strongest known markers. Running SPARK-X independently on all 12 samples gave high cross-method rank concordance (*τ* = 0.778), but SPARK-X called far fewer unique SVGs (mean 84 per sample versus 2,802 for FlashS). Gene Ontology enrichment of method-specific detections showed that FlashS-unique SVGs converged on DNA-binding transcription factor activity in 7 of 12 samples, consistent with layer-specific transcriptional regulation, whereas SPARK-X-unique SVGs yielded no enriched GO terms in 11 of 12 samples (Fig. 4g,h; Supplementary Table 8).

As a stringent cross-platform test, we compared SVG rankings produced independently on mouse brain datasets profiled by Visium (2,702 spots), MERFISH (3.94 million cells), and Slide-seq V2 (41,786 beads). Among 500 genes tested on both Visium and MERFISH, FlashS rankings show significant concordance (*τ* = 0.446, *ρ* = 0.612, permutation *P* < 0.001; Fig. 4d), comparable to PreTSA (*τ* = 0.460); the similar concordance across methods indicates that the strongest spatial signals replicate across platforms regardless of the detection approach. All three pairwise comparisons significantly exceed the permutation null (*P* < 0.001; Fig. 4e,f), with known brain region markers consistently ranked among the top SVGs across all three technologies (Supplementary Fig. 7), indicating that the detected spatial signals reflect genuine biology despite order-of-magnitude differences in cell count and measurement modality.

### Pathway-level recovery in human heart

Cortical layer recovery shows that FlashS retains known brain biology; human heart provides a distinct test of whether the predicted sensitivity gap against fixed smooth bases reveals coherent tissue programs. We compared FlashS, SPARK-X, and PreTSA on a 10x Visium human heart dataset (4,235 spots, 16,716 genes; Fig. 5a). Rank agreement was highest between FlashS and SPARK-X (Kendall *τ* = 0.52, top-50 overlap 80%) and lower against PreTSA (*τ* = 0.41 and 0.38), matching the non-parametric versus parametric divide. Among genes tested by both methods, FlashS detected 3,699 SVGs at *q* < 0.05 compared with 1,123 for PreTSA; 1,030 genes were shared, 2,669 were FlashS-unique, and 93 were PreTSA-unique. Shared SVGs were strongly enriched for cardiac energy metabolism, led by oxidative phosphorylation (*p* < 10^−72^), while FlashS-unique SVGs were enriched for mitochondrial translation and intracellular transport (Fig. 5d; Supplementary Table 5). The PreTSA-unique set yielded no significant GO terms, and visual inspection showed sparse isolated foci rather than broad cardiac tissue patterns (Fig. 5a; Supplementary Fig. 8).

**Fig. 5.**
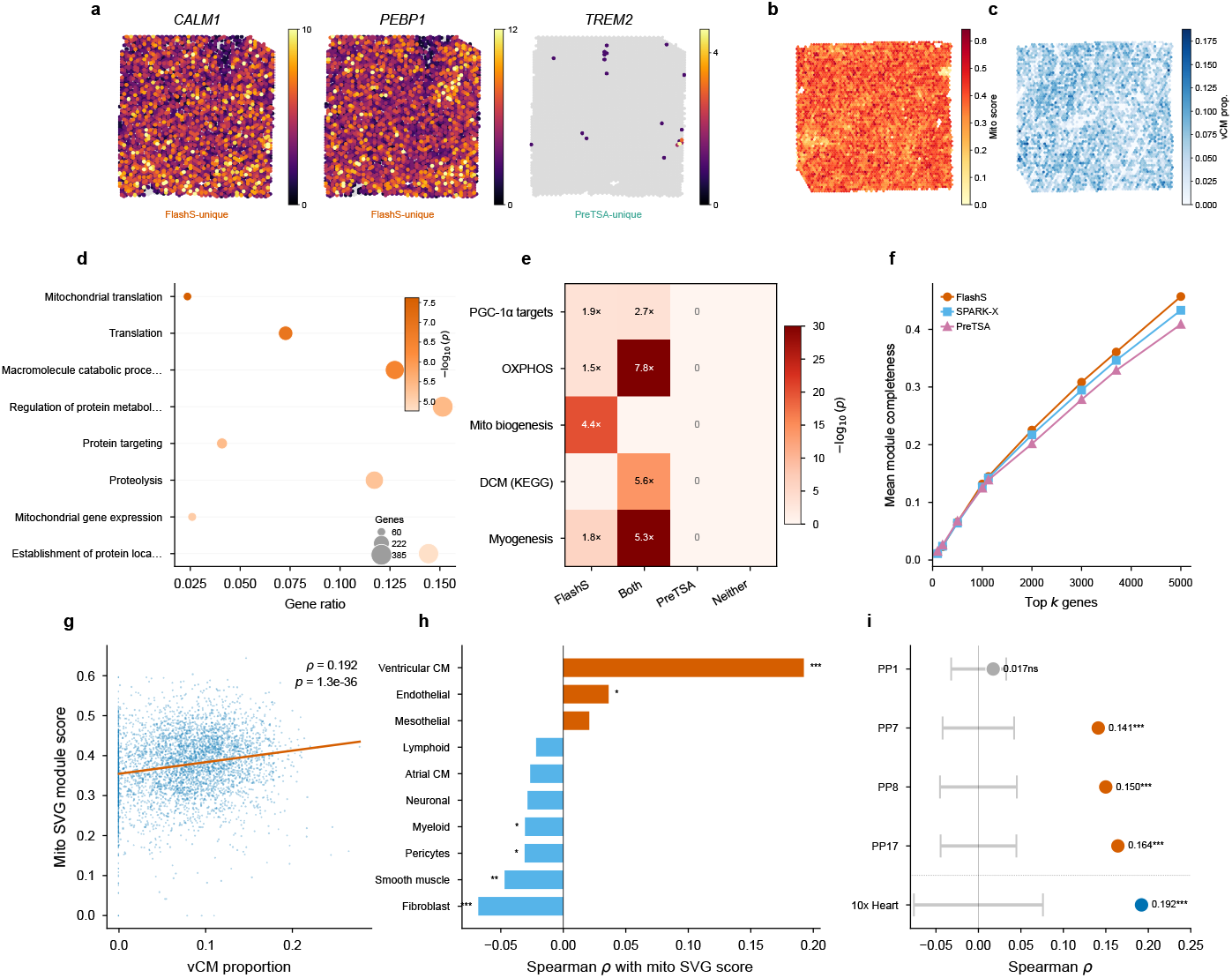
Biological validation and pathway-level coherence in human heart. **a**, Spatial expression of *CALM1* and *PEBP1* (FlashS-unique) and *TREM2* (PreTSA-unique) across 4,235 Visium spots. Color is raw count; gray marks zero. **b**, Mean log-normalized expression of 49 mitochondrial-biogenesis genes across one biologically independent sample. **c**, Deconvolved ventricular-cardiomyocyte (vCM) proportion. **d**, Gene Ontology enrichment of 2,669 FlashS-unique SVGs by a one-sided g:Profiler over-representation test with a 14,634-gene background and g:SCS correction. **e**, One-sided hyper-geometric enrichment of four SVG categories in five MSigDB sets. Benjamini–Hochberg correction was applied across 20 tests. **f**, Mean top-*k* pathway completeness across 365 pathways; distributions are in Supplementary Fig. 10. **g**, Spearman association of vCM proportion and module score across *n* = 4,235 spots (*ρ* = 0.192, exact two-sided *p* = 1.28 *×* 10^−36^). **h**, Spearman associations with cell-type proportions across the same spots. Exact two-sided, unadjusted p-values are 1.28 *×* 10^−36^ (vCM), 0.08495 (atrial cardiomyocyte), 0.01829 (endothelial), 9.44 *×* 10^−6^ (fibroblast), 0.15658 (lymphoid), 0.17621 (mesothelial), 0.04603 (myeloid), 0.06223 (neuronal), 0.04420 (pericytes), and 0.00224 (smooth muscle). **i**, Replication in four biologically independent Kuppe et al.[22] slides. Points are observed *ρ* values; gray intervals span the 2.5th–97.5th percentiles of 10,000 block permutations. Exact two-sided Spearman p-values are 0.25553 (P1), 1.68 *×* 10^−14^ (P7), 8.09 *×* 10^−14^ (P8), 8.24 *×* 10^−14^ (P17), and 1.28 *×* 10^−36^ (discovery). Corresponding 8-by-8 block-permutation p-values are 0.29557, 9.999 *×* 10^−5^, 9.999 *×* 10^−5^, and 9.999 *×* 10^−5^.

To test this, we performed hypergeometric enrichment analysis against five curated gene sets from MSigDB[17], applying Benjamini–Hochberg correction across all 20 tests (5 gene sets × 4 SVG categories; Supplementary Table 7). FlashS-unique SVGs are significantly enriched for PGC-1*α* transcriptional targets (1.90-fold, odds ratio [OR] = 2.39, 95% CI 1.51–3.83, *q* = 1.0 × 10^−3^; Fig. 5e), the master regulator of mitochondrial biogenesis in cardiomyocytes[18, 19]. To test pathway-level recovery we assembled a set of 49 canonical mitochondrial-biogenesis components (full composition and gene list in Methods and Supplementary Table 11). Of these, 40 are detected as SVGs by FlashS—39 appearing exclusively in the FlashS-only category (4.36-fold, *q* = 9.5 × 10^−20^)—with no overlap in the PreTSA-only category. This sensitivity advantage extends beyond the parametric–non-parametric divide. Moran’s I, a standard global autocorrelation statistic, detects only 28 of 49 (57%) of these genes at *q* < 0.05 despite calling 51% of the transcriptome spatially variable genome-wide; SPARK-X detects 25 (51%; Supplementary Table 11). The resulting detection gradient—FlashS > Moran’s I (28) > SPARK-X (25) > PreTSA (1)—is consistent with multi-scale kernel methods recovering more of this compound-pattern gene set. These 40 genes form a spatially coordinated transcriptional program: they share a common upstream regulator (PGC-1*α*), are co-expressed within myocardial compartments, and their collective spatial signature—focal enrichment in ventricular regions superimposed on tissue-wide metabolic gradients—exemplifies the compound, multi-scale patterns that a fixed-degree polynomial basis is not designed to capture. This enrichment extends to disease-relevant gene sets: dilated cardiomyopathy genes (KEGG; OR = 8.81, 95% CI 5.50–14.28, *q* = 4.6 × 10^−14^ in the shared SVG category) and myogenesis hallmark genes (OR = 8.32, 95% CI 6.12–11.38, *q* = 2.4 × 10^−30^) are enriched among SVGs detected by FlashS, while PreTSA-unique SVGs show no significant overlap with any tested pathway (0 of 20 tests significant after FDR correction; Fig. 5e).

To validate the spatial pattern of these mitochondrial SVGs independently, we performed spatial deconvolution using the Litviňuková et al. human heart single-cell atlas[20] (486,134 cells and nuclei spanning 11 major cardiac cell types; 452,136 retained after our preprocessing) via FlashDeconv[21]. We then computed a per-spot mitochondrial SVG module score (mean log-normalized expression of the 49 curated mitochondrial biogenesis genes) and correlated it with deconvolved cell-type proportions across 4,235 Visium spots (Fig. 5b,c). The mitochondrial SVG score is positively correlated with ventricular cardiomyocyte proportion (Spearman *ρ* = 0.192, *R*^2^ ≈ 3.7%, *p* = 1.3 × 10^−36^; Fig. 5g), consistent with the known high mitochondrial content of ventricular cardiomyocytes[20]. The modest effect size indicates that the mito–cardiomyocyte co-localization is one component of a spatially complex tissue, not a dominant signal. By contrast, atrial cardiomyocyte proportion shows no significant correlation (*ρ* = −0.026, *p* = 0.085), reflecting the lower mitochondrial density in atrial tissue, while fibroblasts are negatively correlated (*ρ* = −0.068, *p* = 9.4 × 10^−6^; Fig. 5h). Among all 12 cell types tested, ventricular cardiomyocytes show the clearest positive association with the mitochondrial SVG pattern.

Two technical controls confirm that this association is genuine rather than circular. Spatial block permutation preserves the vCM–mito correlation (*p*_perm_ < 10^−4^ across three grid resolutions), and re-running FlashDeconv after removing all 49 mitochondrial biogenesis genes from its reference leaves the correlation virtually unchanged (Δ*ρ* = 0.006; Supplementary Fig. 8), excluding gene-set leakage.

A complementary decomposition resolves the mito program into cell-type-composition and within-tissue components: regressing out all 12 deconvolved cell-type proportions prior to spatial testing retains 8 of the 49 mitochondrial biogenesis genes as SVGs (genome-wide, 311 of 3,968 SVGs remain). The dominant signal is therefore tissue-scale and tracks the spatial distribution of ventricular cardiomyocytes, while a finer-scale within-cell-type component persists even after composition is removed— consistent with the compound, multi-scale geometry that motivates the kernel in the first place.

To test whether this mito–cardiomyocyte association generalizes beyond a single dataset, we repeated the full analysis on four healthy control Visium slides from an independent cohort[22] (2,043–4,269 spots per slide). Despite only 16 of the 49 mito biogenesis genes being present in the Kuppe Visium panel, the *SVG-detection layer replicates in all four samples*: FlashS-unique SVGs are significantly enriched for mitochondrial biogenesis in every slide (fold enrichment 2.0–4.0, hypergeometric *p* < 0.02; Supplementary Fig. 8), with the largest slide (P1, 4,269 spots) yielding the strongest enrichment (13*/*16 panel genes detected, fold = 2.55, *p* = 6.9 × 10^−5^). The downstream cell2location association replicates in 3 of 4 samples (P7, P8, P17; *ρ* = 0.14–0.16, *R*^2^ ≈ 2–2.6%, block permutation *p*_perm_ < 10^−4^; Fig. 5i). In P1 the cardiomyocyte association is not significant (*ρ* = 0.017); the module score instead associates with pericytes and endothelial cells (*ρ* ≈ 0.06). This slide-level variability may reflect differences in tissue composition or deconvolution accuracy across slides (full per-sample correlations in Supplementary Fig. 8).

These results are consistent with FlashS recovering a biologically plausible spatial program—PGC-1*α*-driven mitochondrial biogenesis genes that co-localize with ventricular cardiomyocytes—that replicates in the majority of independent samples. The effect sizes are modest (*R*^2^ ≈ 2–4%), as expected for a single pathway within a complex tissue, and the P1 slide does not replicate, so the association should be interpreted as suggestive rather than definitive. Rankings are moderately stable after library-size normalization (Kendall *τ* = 0.75; Supplementary Table 14), indicating that a component of the spatial signal co-varies with total counts, which in cardiac tissue reflects both biological (cell-type composition) and technical (sequencing depth) sources.

Is this pathway-level sensitivity specific to the mitochondrial program, or does it generalize? To answer this we scored *module completeness*—the fraction of a curated gene set detected as SVGs at *q* < 0.05—for each method across 365 pathways (50 MSigDB Hallmark and 315 KEGG). Because FlashS detects more SVGs overall (3,699 versus 2,264 for SPARK-X and 1,123 for PreTSA), a fair comparison controls for call volume. At matched call counts (top-*k* genes ranked by each method’s spatial score; Fig. 5f), all three methods achieve comparable module completeness for the highest-ranked genes (*k* ≤ 500), but FlashS pulls ahead at larger *k*: at *k* = 3,699, FlashS covers 36.1% of pathway genes versus 34.6% for SPARK-X and 32.9% for PreTSA. This indicates that FlashS’s extended detections are enriched for pathway-relevant spatial genes rather than for noise.

### Atlas-scale analysis of whole mouse brain MERFISH

Spatial transcriptomics atlases now routinely profile millions of cells, yet it remains unclear which SVG detection methods can operate at this scale with calibrated inference and informative gene rankings. We applied FlashS and four competing methods to the Allen Brain Cell Atlas MERFISH dataset[23] (3,938,808 matched cells from a 500-gene panel plus 50 blank controls spanning the entire adult mouse brain; Fig. 6).

**Fig. 6.**
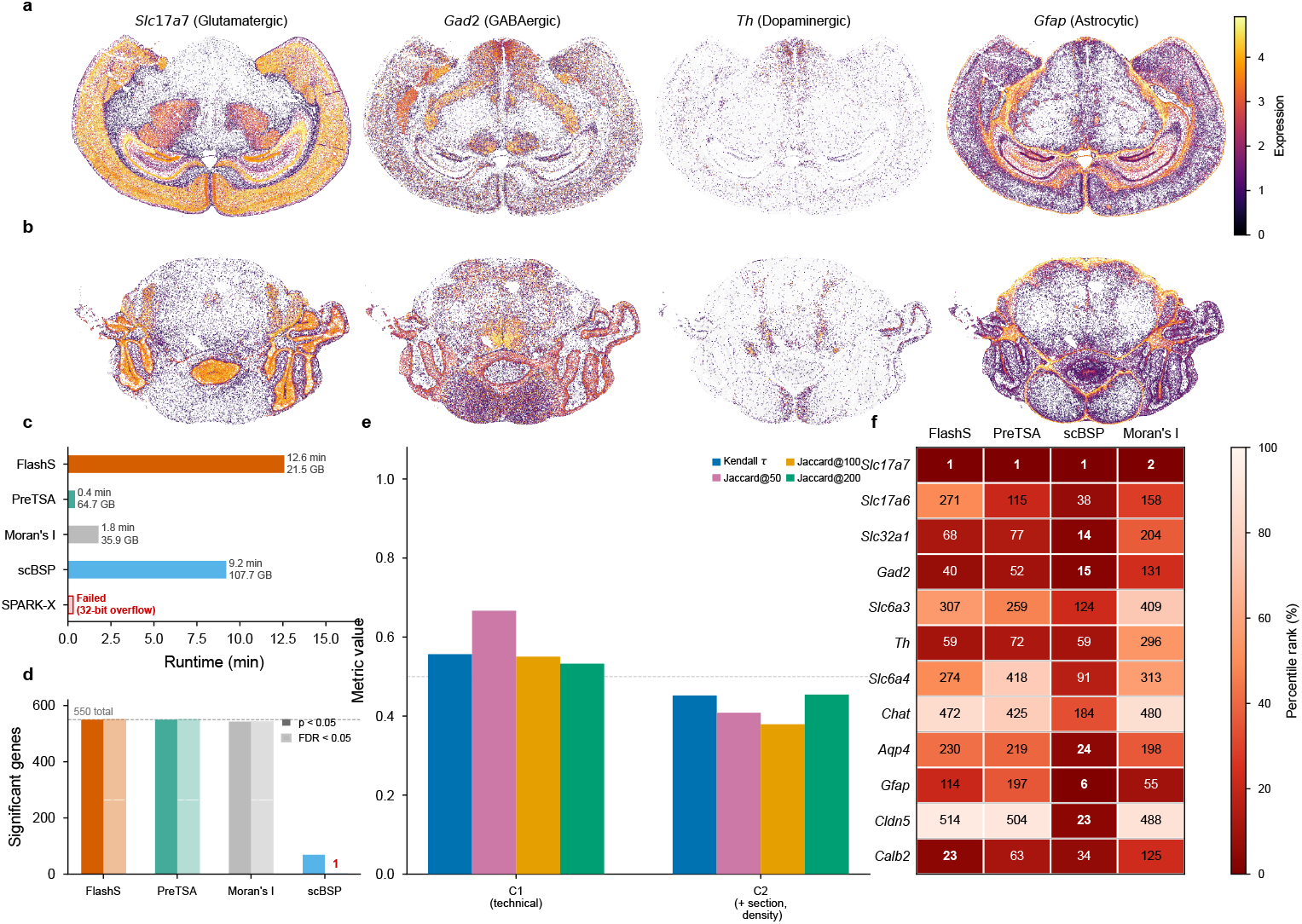
Atlas-scale SVG detection on the Allen MERFISH whole mouse brain. **a,b**, Spatial expression of four representative marker genes in mid-brain (**a**, 115,779 cells) and anterior (**b**) coronal sections: *Slc17a7* (glutamatergic, cortex), *Gad2* (GABAergic), *Th* (dopaminergic, midbrain), and *Gfap* (astrocytic). Color intensity indicates log-transformed expression; gray dots mark non-expressing cells. **c**, Method scalability comparison on 3.94 million cells. PreTSA completes in 25 s with 64.7 GB; SPARK-X fails due to R’s 32-bit integer limit; scBSP requires 108 GB memory. **d**, Number of significant genes detected by each method among 550 tested features at unadjusted *p* < 0.05 and Benjamini–Hochberg *q* < 0.05. P-values are from each method’s native upper-tail omnibus test (the FlashS quadratic-form test, PreTSA F-test, Moran’s I analytical test, or scBSP test); Benjamini– Hochberg correction was applied separately within method across all 550 features. Exact gene-level p- and q-values underlying the counts are provided in Supplementary Data 1. **e**, Robustness of SVG rankings to technical covariate adjustment. Bars show Kendall *τ* (versus unadjusted baseline) and top-*k* Jaccard similarity (*k* = 50, 100, 200) after adjusting for log library size (C1) and section fixed effects with local cell density (C2). **f**, Rankings of 12 prespecified neurotransmitter-system markers across methods (PreTSA ranked by F-statistic to break p-value ties). This panel is descriptive and no hypothesis test was applied. Lower rank indicates stronger spatial signal.

FlashS completed in 12.6 minutes with 21.5 GB peak memory, identifying all 550 matrix features—including the 50 blank negative-control barcodes—as statistically significant (*q* < 0.05). At *n* = 3.94 million cells, p-value saturation is expected for any kernel-based test, making the spatial effect size the meaningful discriminator. Indeed, the 50 blanks cluster at the bottom of the effect-size ranking (median rank 505.5 of 550, 96% in the bottom half; Mann–Whitney *U* test, *p* = 6.2 × 10^−20^; Supplementary Fig. 12), indicating that the effect-size ranking distinguishes real genes from negative controls even when p-values cannot. One outlier (Blank-34, rank 157) may reflect spatially structured technical noise; the remaining 49 blanks have a median rank of 508. Cross-method comparison shows consistent blank demotion across all four methods tested (Supplementary Fig. 12c). Spatial expression maps of representative markers show the known regional distributions (Fig. 6a,b).

The practical advantage of FlashS at this scale emerges from three comparisons. First, *completability* : SPARK-X failed entirely due to R’s 32-bit integer indexing limit; scBSP completed but required 108 GB peak memory—five times that of FlashS—and detected only a single gene after FDR correction (Fig. 6c,d). Second, *memory efficiency* : PreTSA completed in 25 seconds but consumed 64.7 GB, already exceeding standard workstation limits on this 550-gene panel and making whole-transcriptome analysis infeasible at this cell count. Third, *ranking resolution*: at this sample size, p-values saturate near machine precision for all methods; FlashS’s spatial effect size provides a continuous ranking metric that resolves all 550 genes, whereas PreTSA’s p-values yield only 23 distinct values (Fig. 6f). Examination of 12 established neurotransmitter system markers across 8 cell populations confirms that FlashS rankings reflect known spatial biology, with *Slc17a7* (glutamatergic) ranked first (see Methods for full marker list). We assessed the robustness of these rankings to technical covariates by regressing out log library size (C1) and section fixed effects with local cell density (C2) prior to spatial testing. Technical adjustment preserved the overall scale of the effect sizes (median attenuation ratio 1.11; Fig. 6e) but produced moderate reordering of gene ranks (C1: *τ* = 0.56; C2: *τ* = 0.45), indicating that many high-signal genes remain robust while a subset is meaningfully re-ranked. Stratified analyses within individual brain sections (mean *τ* = 0.53) and within single cell types (mean *τ* = 0.41; Supplementary Fig. 6) show that spatial patterns persist within homogeneous subpopulations.

## Discussion

This study supports a direct principle for spatial transcriptomics analysis: preserving a richer spatial basis at scale can reveal biology that restricted bases miss. On controlled simulations, FlashS retains power across hotspot, gradient, domain, and multiscale patterns while maintaining calibrated false-positive rates. On benchmarks, it achieves the highest SVG ranking accuracy across 50 datasets and 9 platforms. On real tissues, its additional detections are enriched for interpretable programs—DNA-binding transcription factor activity in cortex and mitochondrial biogenesis in heart—rather than for isolated genes. The cardiac program is supported by PGC-1*α* target enrichment, ventricular cardiomyocyte co-localization via deconvolution (*R*^2^ ≈ 3.7%), and partial replication in an independent cohort (3 of 4 slides).

The computational result is equally important for current spatial atlases. At million-cell scale, memory rather than wall-clock time often determines whether a method can be used at all. FlashS keeps per-gene working memory at *O*(*D*), independent of the number of spatial locations, and therefore analyses the Allen MERFISH atlas on workstation-class hardware. PreTSA remains faster in wall-clock time because it uses shared dense matrix operations, but its memory scales with the dense expression matrix and becomes limiting for whole-transcriptome atlas analysis. When p-values saturate at very large sample size, FlashS’s spatial effect size provides a continuous ranking that remains useful for prioritizing genes and pathways.

The detection of domain boundaries by a smooth Gaussian kernel (Fig. 3d) warrants a mechanistic note. A single Gaussian kernel cannot represent a sharp spatial boundary, but the multi-scale bandwidth mixture allows local-scale kernels to capture the boundary edge. A hurdle simulation in which spatial signal resides exclusively in the detection probability (zero/nonzero pattern) confirms that FlashS achieves 100% detection power on such patterns, while intensity-only signal at matched strength yields only ∼ 34% TPR. This validates the three-channel design: the binary channel captures spatial information in the zero pattern that rank and direct channels cannot efficiently detect on their own.

Several limitations remain. The Open Problems benchmark is based partly on Matérn Gaussian process simulations, so benchmarks containing sharper boundaries, compositional gradients, and technology-specific noise would be valuable. The kurtosis-corrected two-moment null is calibrated in the settings tested here but cannot represent all higher-order tail behaviour of zero-inflated projections; saddlepoint or resampling-based refinements may improve extreme-tail accuracy at additional cost. The feature count *D* can be reduced to trade accuracy for speed: on the MERFISH atlas, *D* = 50 retains *τ* = 0.90 relative to *D* = 500 at 3.4× speedup (Supplementary Table 15), allowing users to match runtime constraints with graceful degradation.

This work focuses on marginal SVG detection. Conditional tests that adjust for spatial covariates, cell-type composition, or histological features are a natural extension: the Frisch–Waugh–Lovell residualization demonstrated for confounding adjustment on MERFISH (Fig. 6e) can be applied to any set of covariates before the RFF test, extending the same frequency-domain idea from discovery to context-specific spatial modelling. As spatial atlases increasingly provide null reference tissues alongside disease samples, differential SVG testing across conditions will become a key application where the multi-scale flexibility of FlashS may offer particular advantages.

## Conclusions

FlashS recasts SVG detection as frequency-domain spatial testing: random Fourier features preserve Gaussian-kernel flexibility, sparse sketching makes zero-inflated counts computationally favourable, and a kurtosis-corrected null provides analytic calibration without permutation. This combination links scale to biological interpretability: the method analyses a whole-brain MERFISH atlas in minutes—with effect-size rankings that correctly demote negative controls—and identifies a PGC-1*α*-regulated mitochondrial biogenesis program in human heart that maps to ventricular cardiomyocytes and partially replicates in an independent cohort. As spatial atlases grow toward whole organs and organisms, frequency-domain kernel testing offers a practical route to detecting the multi-scale gene programs that organize tissue function.

## Methods

### The FlashS test statistic

To test for spatial variation without constructing *n* × *n* distance matrices, FlashS employs Random Fourier Features (RFF)[9]. By Bochner’s theorem, shift-invariant positive definite kernels can be represented as Fourier transforms of probability measures (Supplementary Note 1). For each spatial location **s**_*i*_ ∈ ℝ^2^, we construct a *D*-dimensional feature vector:

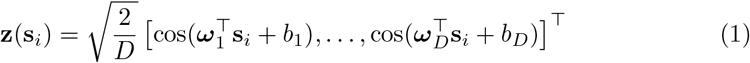

where ***ω***_*k*_ ∼ *N*(0, *σ*^−2^**I**) are sampled from the spectral density of a Gaussian kernel with bandwidth *σ*, and *b*_*k*_ ∼ Uniform(0, 2*π*) are phase offsets. The inner product **z**(**s**), **z**(**s**^*′*^) approximates *k*(**s, s**^*′*^) = exp(−∥**s s**^*′*^ ∥ ^2^*/*2*σ*^2^).

#### Adaptive multi-scale bandwidth

Because spatial expression spans scales from cellular neighborhoods to tissue-wide gradients, FlashS sets a local scale from the median nearest-neighbor distance and a global scale from the Fiedler value[11] *λ*_2_ of the graph Laplacian, giving 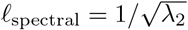 bandwidths (default *L* = 7) are log-spaced between these two scales, and the *D* features are distributed across scales by divmod(*D, L*).

#### Sparse sketching

For each gene, FlashS tests whether standardized expression **y** associates with **Z** via 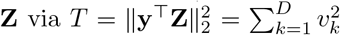. The projections *v*_*k*_ are computed without materializing **Z**[24] (Supplementary Note 2):

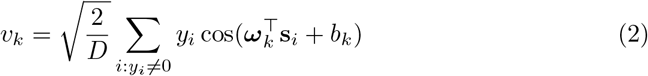

achieving *O*(nnz ·*D*) per gene. Centering is implicit: the column sums **1**^⊤^**Z** are pre-computed once in one exact *O*(*nD*) pass (with *O*(*D*) memory) and subtracted as 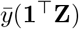, preserving sparsity. Subsampling the centering vector is not viable—centering error enters the quadratic statistic as 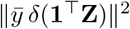 and dominates the null scale when *n* is much larger than the moment-estimation subsample.

#### Three-part test for zero-inflated data

Three complementary channels are applied per gene and bandwidth: a *binary* test on **1**(*y* > 0) (detection pattern), a *rank* test on rank-transformed non-zero values (outlier-robust intensity), and a *direct* test on raw expression (preserves library-size-correlated signal). A linear spatial trend test for each channel—regressing the channel vector onto centered coordinates—supplements the RFF kernels to capture tissue-wide gradients that fall between Gaussian scales.

#### Cauchy combination

Per-scale and per-channel p-values are combined[12] as

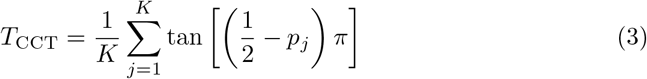

with *K* = 3*L* + 3 (three channels across *L* scales plus three linear trend tests). Under the null, *T*_CCT_ follows a standard Cauchy distribution regardless of dependency among component tests, yielding a single combined p-value per gene.

### Null distribution and multiple testing

Under the null, the centered projections are *v*_*k*_ = **y** ^⊤^**z**_*k*_ and 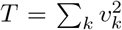. The RFF columns are correlated—particularly at large bandwidths where multiple cosine features approximate similar low-frequency modes—so the standard Satterthwaite approximation assuming independent *v*_*k*_ underestimates the null variance. For jointly normal *v*_*k*_, 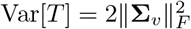, with 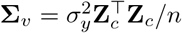 and **Z**_*c*_ the column-centered feature matrix.

We use a kurtosis-corrected Satterthwaite moment match[25], with 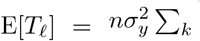 Var [*z*_*k,l*_] and per-scale variance

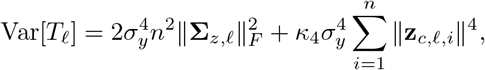

where **Σ**_*z,l*_ is the centered feature covariance within scale *l* and 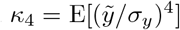 −3 is the per-gene excess kurtosis[26]. The kurtosis term—absent from the quadratic-form approximations used by existing SVG methods—corrects for the leptokurtic distribution of zero-inflated expression (*κ*_4_ ≫ 0), which otherwise inflates p-values (Supplementary Note 3). Both terms are estimated once on a coordinate subsample (*M* = 10,000 by default) at 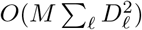 cost, yielding *κ* = Var[*T*_*l*_]*/*(2 E[*T*_*l*_]) and *v* = 2 E[*T*_*l*_]^2^*/*Var[*T*_*l*_] for a 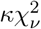 reference. This provides analytic p-values without permutation. Gene-level p-values are adjusted by Benjamini–Hochberg at level *α* = 0.05; the same correction is applied to every compared method.

#### Open Problems benchmark

We evaluated FlashS on the Open Problems SVG benchmark[13] (50 datasets, 11 leaderboard methods; benchmark snapshot archived at tag `flashs-benchmark-v1`^1^, prior to the 2025 expansion to 96 datasets). Methods are scored with default parameters, and Kendall *τ* is computed between each method’s per-gene spatial score and the benchmark’s ground truth. As in Open Problems, each method uses the score most appropriate to its model: SpatialDE, SpatialDE2, and SOMDE report the fraction of spatial variance (FSV); Moran’s I reports *I*; SpaGFT and Spanve report method-specific statistics; scGCO uses −log_10_(*p*). FlashS uses the spatial effect size—the maximum ratio of observed to expected test statistic across the three test channels—because an ablation across all 50 datasets shows that − log_10_(*p*) alone yields substantially lower concordance (*τ* = 0.868) as p-values saturate at moderate *n*, whereas the effect size provides a continuous, non-saturating ranking (Supplementary Table 13).

FlashS was run on raw counts with *D* = 500, *L* = 7, and the three-part test. Three methods not in the original leaderboard—PreTSA[8], Hotspot[7], and scBSP[6]—were benchmarked on the same 50 datasets with default parameters; PreTSA was run with its default knot setting (cubic B-spline, no internal knots) on library-size-normalized and log-transformed data as the authors recommend. All benchmarks were performed on 16-core AMD EPYC compute nodes. The reported analyses were performed with FlashS v0.1.0 using Python 3.11, NumPy 1.26, SciPy 1.12, and Numba 0.59. The Zenodo archive contains the manuscript-associated v0.2.0 release together with the analysis scripts; v0.1.0 is retained here to identify the exact software version used to generate the reported results.

### Scalability experiments

Scalability was evaluated on simulated datasets with 5,000 genes and cell counts from 10^3^ to 10^6^ (3 replicates per condition). Spatial coordinates were uniform; expression was negative binomial (*n* = 1, *p* = 0.1; ∼90% zeros). Runtime was measured from model fitting through p-value computation; peak memory was recorded via the `resource` module. Runtime comparison against Moran’s I (Squidpy[27]), SpatialDE[1], Hotspot[7], SPARK-X[5], SOMDE[28], scBSP[6], Spanve[29], nnSVG[4], and PreTSA[8] was performed up to 5 × 10^4^ cells (3 replicates; 2-hour timeout, 50 GB memory limit). SPARK-X and nnSVG were called via rpy2. Extended FlashS–PreTSA comparison was performed up to 10^6^ cells (Supplementary Table 4). All experiments used 16-core AMD EPYC compute nodes.

### Simulation studies

#### Calibration

Null datasets on a 50 × 50 grid (2,500 spots) at three zero-fraction levels— ∼50% (Poisson, *λ* = 0.7), ∼90% (negative binomial, *n* = 1, *p* = 0.9), and ∼95% (negative binomial with additional 50% dropout)—were generated without spatial structure (20 replicates of 1,000 genes each, 60,000 null tests).

#### Power and pattern benchmarks

On the same grid with 500 SVGs and 500 nulls per replicate (10 replicates), we evaluated hotspot (localized Gaussian envelope), gradient (linear ramp), and periodic (sinusoidal) patterns at effect sizes 0.1–0.5 and sparsity 10–50%. For cross-method power comparison (Fig. 3d–f), four pattern types— gradient, hotspot, domain boundary (sharp step), and multiscale (gradient + hotspot)— were simulated with 200 SVGs and 200 nulls per replicate (10 replicates), effect sizes {0.3, 0.5, 1.0}, and sparsity {50%, 80%, 95%}; background was negative binomial. FlashS, SPARK-X, and PreTSA were applied with default parameters.

#### Ablations

*D* ∈ {50, 100, 200, 500, 1,000} with *L* = 7 fixed, and *L* ∈ {1, 3, 5, 7, 10} with *D* = 500 fixed (5 replicates per condition).

### Atlas-scale MERFISH analysis

FlashS, PreTSA, SPARK-X, scBSP, and Moran’s I (Squidpy) were applied to the Allen Brain Cell Atlas MERFISH dataset[23] (specimen C57BL6J-638850). The processed expression matrix contains 4,334,174 cells ×550 features (500-gene panel + 50 blank controls); after matching cell identifiers between the H5AD expression matrix and spatial metadata, 3,938,808 cells were retained. Expression was used as raw counts. FlashS ran with *D* = 500, *L* = 7; SPARK-X via rpy2; scBSP (v0.3.1) via its Python interface; Moran’s I via Squidpy (*k* = 6); PreTSA on library-size-normalized and log-transformed data (rankings from the F-statistic, since p-values saturate at machine zero at this *n*). Each method was an isolated job on a 16-core compute node with 128 GB memory. Biological validation used 12 established neurotransmitter/glial markers: *Slc17a7, Slc17a6* (glutamatergic); *Slc32a1, Gad2* (GABAergic); *Slc6a3, Th* (dopaminergic); *Slc6a4* (serotonergic); *Chat* (cholinergic); *Aqp4, Gfap* (astrocytic); *Cldn5* (endothelial); and *Calb2* (interneuron).

#### Structured null testing

Three exchangeability strategies[30] were applied to the full atlas: (i) global row permutation (N0; shuffles across all cells, destroying all spatial structure), (ii) within-section permutation (N1; preserves between-section structure), and (iii) spatial block permutation (N2; 6 × 6 grid per section, preserves within-block autocorrelation). Replicate counts: N0 = 3 (1,650 null tests), N1 = 10 (5,500 tests), N2 = 6 (3,300 tests). Rejection rate was the fraction of genes with *p* < 0.05; for N0 this is the empirical false positive rate (Supplementary Table 9).

#### Confounding sensitivity

We residualized expression against three covariate sets— C0 (no adjustment), C1 (log total counts; no mitochondrial genes are present in the 550-gene panel), C2 (C1 + brain-section fixed effects + local cell density from a 50 × 50 2D histogram per section)—using Frisch–Waugh–Lovell demeaning for the categorical fixed effect followed by OLS projection for continuous covariates. FlashS was then applied with `min expressed` = 0 (residuals are not sparse). Stability was assessed by Kendall *τ* and top-*k* Jaccard against C0. Stratified analyses ran FlashS within the 5 largest brain sections and the 5 most abundant cell-type classes.

### Real-data validation and biological enrichment

#### DLPFC cross-sample consistency

FlashS and SPARK-X were run on 12 Visium samples from the Human DLPFC dataset[16] (3 donors × 4 samples; 33,538 genes per sample). FlashS used defaults on raw counts; SPARK-X ran independently per sample via rpy2. Cross-sample consistency was quantified by pairwise Kendall *τ* of effect-size rankings. Marker enrichment was evaluated against 27 cortical layer markers curated from Maynard et al. (L1–L6 + white matter); fold enrichment at top-*k* is the fraction of markers in the top *k* divided by the expected fraction under random ranking.

#### Cross-platform concordance

SVG rankings were compared across three independently profiled mouse brain datasets—10x Visium (2,702 spots, 32,285 genes), Allen MERFISH (3,938,808 cells, 550 genes), and Slide-seq V2 hippocampus (41,786 beads, 4,000 HVGs)—by Kendall *τ*, Spearman *ρ*, and top-*k* Jaccard on common genes, with significance from 1,000-replicate permutation tests.

#### Gene Ontology enrichment

Method-specific SVGs were tested with g:Profiler[31] (GO:BP/MF/CC, KEGG, Reactome; g:SCS *p* < 0.05). Background was the set of genes tested by both methods (14,634 for the heart; per-sample for DLPFC). Genes were stratified by method-specific detection at *q* < 0.05.

#### Heart metabolic validation

Hypergeometric enrichment used five curated gene sets from MSigDB[17]: PPARGC1A TARGET GENES (78 of 83 in universe), HALLMARK OXIDATIVE PHOSPHORYLATION (199), a 49-gene mitochondrial biogenesis set (MRPL/MRPS, TOMM/TIMM, ETC assembly factors; full list in Supplementary Table 11), KEGG DILATED CARDIOMYOPATHY (71), and HALL-MARK MYOGENESIS (177). Spatial deconvolution used the Litviňuková single-cell atlas[20] (486,134 cells; 452,136 retained across 12 cell types) via FlashDeconv[21] with defaults (sketch dimension 512, 2,000 HVGs). A per-spot mito-module score (mean log_1p_-normalized expression of the 49 genes) was correlated with deconvolved cell-type proportions via Spearman rank correlation; Moran’s I (Squidpy, *k* = 6) and SPARK-X were run for comparison. Spatial block permutation (*m* ∈ {6, 8, 10}; 10,000 permutations) accounted for spatial autocorrelation. Gene-set leakage was excluded by re-running deconvolution after removing the 49 genes from the expression matrix. Module completeness across 365 gene sets (50 MSigDB Hallmark 2020, 315 KEGG 2021 Human) was |*G* ∩ SVG_*M*_ ∩ *U*| */* |*G* ∩ *U* |, excluding sets with fewer than 5 genes in *U* .

#### External cardiac replication

Four healthy control Visium slides from Kuppe et al.[22] (P1, P7, P8, P17; 2,043–4,269 spots per slide, 13,144–15,730 genes after filtering) were analyzed with identical parameters to the discovery cohort. Pre-computed cell2location[32] deconvolution proportions from the source data were used directly; all four slides are from the left ventricle, so cardiomyocyte proportion serves as a ventricular cardiomyocyte proxy. The transferred 49-gene mito-module score and spatial block permutation tests followed the discovery protocol. Only 16 of the 49 biogenesis genes are present in the Kuppe Visium panel.

### Implementation

FlashS is implemented in Python with NumPy/SciPy and Numba[33]-compiled parallel kernels. Key optimizations include sparse sketching for *O*(nnz · *D*) projection (full per-gene cost *O*(nnz · *D* + nnz log nnz) including rank transformation), implicit centering to preserve sparsity, and a fused triple kernel that evaluates all three test channels in a single pass over the non-zero entries. Model fitting uses one exact *O*(*nD*) centering pass plus subsampled moment estimation; this keeps inference calibrated while preserving scalability in the gene-wise testing stage. Default parameters: *D* = 500, *L* = 7. Source code: https://github.com/cafferychen777/FlashS.

### Statistics and Reproducibility

All statistical tests, sample sizes, replicate definitions, and multiple-testing procedures are specified in the corresponding Methods subsections and figure legends. Null-calibration experiments used *n* = 20 independent simulated datasets per sparsity condition, each containing 1,000 independently generated null genes. Power experiments used *n* = 10 independent simulated datasets per pattern, effect-size, and sparsity condition, each containing 200 spatial and 200 null genes. Computational scaling experiments used *n* = 3 independent process-level runs per method and sample size. Full-atlas null calibration used *n* = 3 independent global permutations of the 550-feature MERFISH matrix. DLPFC validation used 12 biologically independent tissue sections from 3 donors, and cardiac replication used 4 biologically independent healthy-control Visium slides. No data were excluded after analysis, no statistical method was used to predetermine sample size, and analyses were not randomized or blinded because they used public observational datasets or computer-generated simulations. Gene-level p-values were adjusted within each analysis universe by the Benjamini– Hochberg procedure unless otherwise stated. Enrichment tests and spatial permutation tests are described with their sidedness and number of permutations in the relevant Methods and legends. Exact numerical source data underlying all main-text graphs are provided in Supplementary Data 1.

## Supplementary information

Supplementary information is available for this paper.

## Declarations

### Ethics approval and consent to participate

Not applicable. This study uses only previously published, publicly available datasets.

### Consent for publication

Not applicable.

### Authors’ contributions

C.Y., X.Z., and J.C. jointly conceptualised the study. C.Y. developed the methodology, implemented the computational framework, conducted formal analysis, and performed data curation, visualisation, and validation. X.Z. and J.C. jointly supervised the research, secured funding, provided resources, served as corresponding authors, and administered the project, contributing equally in these senior roles. All authors participated in manuscript writing and revision, and read and approved the final manuscript.

## Acknowledgements

We thank the Open Problems benchmark suite, the Allen

Brain Cell Atlas, and 10x Genomics for making their datasets publicly available, and the developers of the open-source tools used in this work. Portions of this research were conducted with the advanced computing resources provided by the Texas A&M Department of Statistics Arseven Computing Cluster.

## Data availability

All datasets analysed during this study are publicly available from their original sources. The Allen Brain Cell Atlas MERFISH data are available in the Brain Knowledge Platform repository, https://portal.brain-map.org/atlases-and-data/bkp/abc-atlas. The 10x Visium human heart dataset is available in the 10x Genomics datasets repository, https://www.10xgenomics.com/datasets/human-heart. The human dorsolateral prefrontal cortex Visium data[16] are available in the spatialLIBD Bioconductor repository, http://spatial.libd.org/spatialLIBD/. The Slide-seq V2 hippocampus dataset[34] is available in the Broad Institute Single Cell Portal. The Open Problems SVG benchmark datasets are available through the Open Problems framework[13]. The Kuppe et al. healthy-control Visium slides[22] are available from Zenodo, https://zenodo.org/records/6578047. The Litviňuková et al. human heart single-cell atlas[20] is available at https://www.heartcellatlas.org. Source data underlying all graphs in Figs. 2–6 are provided in Supplementary Data 1.

## Code availability

FlashS v0.2.0 and the scripts used to reproduce the analyses and figures are available under the MIT license at https://github.com/cafferychen777/FlashS. This manuscript-associated release is permanently archived on Zenodo at https://doi.org/10.5281/zenodo.21724351[35]; the reported analyses were generated with v0.1.0, as specified in the Methods. The exact software versions and analysis parameters are reported in the Methods and in the repository environment files.

## List of Figures

## Footnotes

1 https://github.com/cafferychen777/task_spatially_variable_genes/tree/flashs-benchmark-v1

