## Supplementary Information for "FlashS reveals multiscale spatial gene programs at atlas scale"

**4 Supplementary Tables** **10**

**5 Supplementary Figures** **11**

### 1 Supplementary Note 1: Random Fourier Feature Approximation Theory

This note summarizes the Random Fourier Feature (RFF) construction used in FLASHS and derives the multi-scale extension that underlies the FLASHS feature matrix. The Bochner and Monte-Carlo machinery is standard [10]; we state only the facts invoked elsewhere in the supplement.

#### 1.1 RFF construction (Rahimi and Recht, 2007)

By Bochner's theorem, any continuous shift-invariant positive-definite kernel $k$ with $k(0) = 1$ admits the representation $k(\mathbf{s} - \mathbf{s}') = \mathbb{E}_{\omega\sim p}[\cos(\boldsymbol{\omega}^\top(\mathbf{s} - \mathbf{s}'))]$, where the spectral density $p$ is the Fourier transform of $k$. For the Gaussian kernel $k(\mathbf{s}, \mathbf{s}') = \exp(-\|\mathbf{s} - \mathbf{s}'\|^2/(2\sigma^2))$, $p(\boldsymbol{\omega}) = \mathcal{N}(\mathbf{0}, \sigma^{-2}\mathbf{I}_d)$. Introducing an independent phase $b \sim \mathrm{Unif}(0, 2\pi)$ yields $k(\mathbf{s}, \mathbf{s}') = 2\,\mathbb{E}_{\omega,b}[\cos(\boldsymbol{\omega}^\top\mathbf{s} + b)\cos(\boldsymbol{\omega}^\top\mathbf{s}' + b)]$, which becomes the RFF estimator when the expectation is replaced by a Monte Carlo average.

**Definition 1** (Random Fourier Feature map)**.** For $\boldsymbol{\omega}_j \overset{\text{iid}}{\sim} \mathcal{N}(\mathbf{0}, \sigma^{-2}\mathbf{I})$ and $b_j \overset{\text{iid}}{\sim} \mathrm{Unif}(0, 2\pi)$,

$$\mathbf{z}(\mathbf{s}) = \sqrt{\tfrac{2}{D}}\,[\cos(\boldsymbol{\omega}_1^\top\mathbf{s} + b_1),\, \ldots,\, \cos(\boldsymbol{\omega}_D^\top\mathbf{s} + b_D)]^\top \in \mathbb{R}^D. \tag{1}$$

**Theorem 1** (Unbiasedness and pointwise concentration)**.** $\mathbb{E}[\langle\mathbf{z}(\mathbf{s}), \mathbf{z}(\mathbf{s}')\rangle] = k(\mathbf{s}, \mathbf{s}')$*. Since each summand is bounded by* $2/D$*, Hoeffding's inequality gives*

$$\Pr\big[|\langle\mathbf{z}(\mathbf{s}), \mathbf{z}(\mathbf{s}')\rangle - k(\mathbf{s}, \mathbf{s}')| > \epsilon\big] \le 2\exp\Big(-\tfrac{D\epsilon^2}{2}\Big)\,. \tag{2}$$

**Proposition 2** (Pointwise RFF bound)**.** *Equation* (2) *holds for every fixed pair* $(\mathbf{s}, \mathbf{s}')$*. Setting* $\epsilon = 0.12$ *and* $\delta = 0.05$ *yields* $D \ge 500$*, matching the* FLASHS *default.*

Because the test statistic aggregates across $D$ projections and $L$ scales, per-evaluation approximation error averages out: the ablation in Supplementary Fig. 1a–b shows that $D = 500$ reproduces benchmark accuracy within 0.2% of $D = 1{,}000$.

#### 1.2 Extension to Multi-scale Kernels

Biological spatial patterns occur at multiple characteristic scales—from fine-grained cellular neighborhoods to tissue-wide gradients. A single bandwidth $\sigma$ may miss patterns at other scales. FLASHS addresses this by constructing a multi-scale feature map.

Given $L$ bandwidth scales $\sigma_1 < \sigma_2 < \cdots < \sigma_L$, we construct:

$$\mathbf{Z} = \begin{bmatrix}\mathbf{Z}^{(1)} & \mathbf{Z}^{(2)} & \cdots & \mathbf{Z}^{(L)}\end{bmatrix} \in \mathbb{R}^{n\times D} \tag{3}$$

where $\mathbf{Z}^{(m)} \in \mathbb{R}^{n\times D_m}$ contains $D_m$ features sampled with bandwidth $\sigma_m$. Features are distributed via $\mathrm{divmod}(D, L)$: each scale receives $\lfloor D/L\rfloor$ features, with the first $D \bmod L$ scales receiving one extra.

**Proposition 3** (Multi-scale Kernel Approximation)**.** *The concatenated feature map approximates a mixture of Gaussian kernels:*

$$\langle \mathbf{z}(\mathbf{s}), \mathbf{z}(\mathbf{s}') \rangle \approx \sum_{m=1}^{L} w_m \cdot k_{\sigma_m}(\mathbf{s}, \mathbf{s}') \tag{4}$$

*where* $k_{\sigma_m}(\mathbf{s}, \mathbf{s}') = \exp(-\|\mathbf{s} - \mathbf{s}'\|^2 / 2\sigma_m^2)$ *and* $w_m = D_m/D$.

*Proof.* Each block uses the global normalization $\sqrt{2/D}$, so for a block with $D_m$ features sampled at bandwidth $\sigma_m$, Theorem 1 gives $\mathbb{E}[\langle \mathbf{z}^{(m)}(\mathbf{s}), \mathbf{z}^{(m)}(\mathbf{s}') \rangle] = (D_m/D)\, k_{\sigma_m}(\mathbf{s}, \mathbf{s}')$. The inner product of the concatenated features is:

$$\begin{aligned} \langle \mathbf{z}(\mathbf{s}), \mathbf{z}(\mathbf{s}') \rangle &= \sum_{m=1}^{L} \langle \mathbf{z}^{(m)}(\mathbf{s}), \mathbf{z}^{(m)}(\mathbf{s}') \rangle & (5) \\ &= \sum_{m=1}^{L} \frac{2}{D} \sum_{j \in \text{block } m} \cos(\boldsymbol{\omega}_j^\top \mathbf{s} + b_j) \cos(\boldsymbol{\omega}_j^\top \mathbf{s}' + b_j) & (6) \end{aligned}$$

Taking expectations and noting that each block has $D_m$ features with scale $\sqrt{2/D}$, the result follows. □

*Remark* 1. In FlashS, bandwidths are selected adaptively based on spatial geometry: the smallest bandwidth captures local patterns (derived from nearest-neighbor distances), while the largest captures tissue-wide gradients (derived from spatial extent). Intermediate bandwidths are log-spaced between these extremes. This ensures detection of patterns at all biologically relevant scales without manual tuning.

### 2 Supplementary Note 2: Sparse Sketching Error Bounds

This note establishes the theoretical guarantees for the sparse sketching approach used in FlashS for global SVG detection.

#### 2.1 Problem Setup

For global SVG detection, we compute the test statistic:

$$T = \|\mathbf{y}^\top \mathbf{Z}\|_2^2 = \sum_{k=1}^{D} \left( \sum_{i=1}^{n} y_i Z_{ik} \right)^2 \tag{7}$$

where $\mathbf{y} \in \mathbb{R}^n$ is the (standardized) expression vector and $\mathbf{Z} \in \mathbb{R}^{n \times D}$ is the RFF matrix.

#### 2.2 Sparse Computation

For sparse expression data (common in single-cell/spatial transcriptomics), let $\text{nnz} = |\{i : y_i \neq 0\}|$ denote the number of non-zero entries.

The key observation is that we can compute each component:

$$v_k = \sum_{i=1}^{n} y_i Z_{ik} = \sum_{i:y_i \neq 0} y_i Z_{ik} = \sqrt{\frac{2}{D}} \sum_{i:y_i \neq 0} y_i \cos(\boldsymbol{\omega}_k^\top \mathbf{s}_i + b_k) \tag{8}$$

in $O(\text{nnz})$ time without constructing $\mathbf{Z}$ explicitly.

#### 2.3 Computational Complexity

**Proposition 4.** *The test statistic $T = \sum_{k=1}^{D} v_k^2$ can be computed in $O(nnz \cdot D)$ time and $O(D)$ space.*

*Proof.*

1. Pre-compute and store $\{\boldsymbol{\omega}_k, b_k\}_{k=1}^{D}$: $O(dD)$ space
2. For each $k = 1, \ldots, D$:
   - Initialize $v_k = 0$
   - For each $i$ with $y_i \neq 0$: compute $v_k \mathrel{+}= y_i \cos(\boldsymbol{\omega}_k^\top \mathbf{s}_i + b_k)$
   - This takes $O(\text{nnz})$ time per $k$
3. Sum $T = \sum_k v_k^2$: $O(D)$ time

Total: $O(\text{nnz} \cdot D)$ time. Space for $\{v_k\}$: $O(D)$. □

#### 2.4 Implicit Centering: Preserving Sparsity

A critical challenge arises from the requirement that expression vectors be centered (zero mean) for valid kernel-based testing. Naively, centering destroys sparsity: if $\tilde{\mathbf{y}} = \mathbf{y} - \bar{y}\mathbf{1}$ is the centered vector, then $\text{nnz}(\tilde{\mathbf{y}}) = n$ even when $\text{nnz}(\mathbf{y}) \ll n$.

FlashS employs *implicit centering* to maintain sparse projection complexity at $O(\text{nnz} \cdot D)$ per gene. Let $\mathbf{y}$ be the sparse raw expression vector (e.g., counts or ranks) and $\tilde{\mathbf{y}} = \mathbf{y} - \bar{y}\mathbf{1}$ be the centered version where $\bar{y} = \frac{1}{n}\sum_i y_i$.

**Proposition 5** (Implicit Centering)**.** *The projection of centered data can be computed without materializing the dense centered vector:*

$$\tilde{\mathbf{v}} = \tilde{\mathbf{y}}^\top \mathbf{Z} = \mathbf{y}^\top \mathbf{Z} - \bar{y}(\mathbf{1}^\top \mathbf{Z}) \tag{9}$$

*where:*

- $\mathbf{y}^\top \mathbf{Z}$ *is computed in $O(nnz \cdot D)$ using only non-zero entries of* $\mathbf{y}$
- $\mathbf{1}^\top \mathbf{Z}$ *(the column sums of* $\mathbf{Z}$*) is pre-computed once in $O(n \cdot D)$ during model fitting*

*Proof.* By linearity of matrix multiplication:

$$\tilde{\mathbf{v}} = (\mathbf{y} - \bar{y}\mathbf{1})^\top \mathbf{Z} \tag{10}$$

$$= \mathbf{y}^\top \mathbf{Z} - \bar{y} \cdot \mathbf{1}^\top \mathbf{Z} \tag{11}$$

The term $\mathbf{y}^\top \mathbf{Z}$ requires summing over non-zero entries only:

$$[\mathbf{y}^\top \mathbf{Z}]_k = \sum_{i:y_i \neq 0} y_i Z_{ik} \tag{12}$$

which costs $O(\text{nnz})$ per feature, or $O(\text{nnz} \cdot D)$ total.

The term $\mathbf{1}^\top \mathbf{Z} = \sum_{i=1}^n \mathbf{z}(\mathbf{s}_i)^\top$ is independent of $\mathbf{y}$ and is pre-computed during `fit()`, requiring $O(n \cdot D)$ time once for all genes.

Thus, the per-gene complexity remains $O(\text{nnz} \cdot D)$, preserving the sparse efficiency. □

*Remark* 2. This implicit centering technique was pioneered by SPARK-X for scalable SVG detection. FlashS extends this approach with true sketching that avoids storing the $n \times D$ feature matrix entirely.

#### 2.5 Approximation Quality

Let $K \in \mathbb{R}^{n \times n}$ be the true kernel matrix with $K_{ij} = k(\mathbf{s}_i, \mathbf{s}_j)$, and let $\hat{K} = \mathbf{Z}\mathbf{Z}^\top$ be the RFF approximation.

**Theorem 6** (Kernel Matrix Approximation)**.** *For any $\epsilon, \delta > 0$, if $D = O(\epsilon^{-2} \log(n/\delta))$, then with probability at least $1 - \delta$:*

$$\|K - \hat{K}\|_{\max} \leq \epsilon \tag{13}$$

*where $\|\cdot\|_{\max}$ denotes the element-wise maximum absolute value.*

*Proof.* Apply Proposition 2 with a union bound over all $\binom{n}{2} + n \leq n^2$ pairs. □

#### 2.6 Test Statistic Approximation

The true kernel-based statistic is $T^* = \mathbf{y}^\top K \mathbf{y}$, and our approximation is $\hat{T} = \mathbf{y}^\top \hat{K} \mathbf{y} = \|\mathbf{y}^\top \mathbf{Z}\|_2^2$.

**Theorem 7** (Statistic Approximation)**.** *Under the conditions of Theorem 6:*

$$|T^* - \hat{T}| \leq \epsilon \|\mathbf{y}\|_1^2 \tag{14}$$

*with probability at least $1 - \delta$.*

*Proof.*

$$|T^* - \hat{T}| = |\mathbf{y}^\top (K - \hat{K})\mathbf{y}| \tag{15}$$

$$= \left| \sum_{i,j} y_i y_j (K_{ij} - \hat{K}_{ij}) \right| \tag{16}$$

$$\leq \sum_{i,j} |y_i||y_j||K_{ij} - \hat{K}_{ij}| \tag{17}$$

$$\leq \|K - \hat{K}\|_{\max} \left( \sum_i |y_i| \right)^2 \tag{18}$$

$$= \epsilon \|\mathbf{y}\|_1^2 \tag{19}$$

□

*Remark* 3. For standardized expression vectors with $\|\mathbf{y}\|_2 = 1$, we have $\|\mathbf{y}\|_1 \leq \sqrt{n}$, so the bound becomes $|T^* - \hat{T}| \leq \epsilon n$. Since $T^* = O(n)$ under the alternative, the relative error is $O(\epsilon)$.

#### 2.7 Comparison with Dense Methods

| **Method** | **Time** | **Space** | **Error** |
|---|---|---|---|
| Exact kernel ($K$) | $O(n^2)$ | $O(n^2)$ | 0 |
| Dense RFF ($\mathbf{ZZ}^\top$) | $O(nD) + O(n^2)$ | $O(nD)$ | $O(1/\sqrt{D})$ |
| Sparse RFF (ours) | $O(\text{nnz} \cdot D)$ | $O(D)$ | $O(1/\sqrt{D})$ |

For sparse spatial transcriptomics data with $\text{nnz} \ll n$ (e.g., 10% non-zero rate), our approach achieves a 10× speedup over dense RFF while maintaining the same approximation quality. For the full FlashS testing pipeline, per-gene complexity is $O(\text{nnz} \cdot D + \text{nnz} \cdot \log \text{nnz})$, where the additional term comes from rank transformation of non-zero entries.

#### 2.8 Practical Considerations

1. **Choice of $D$**: We use $D = 500$ by default. By Proposition 2, this provides $\epsilon \approx 0.12$ with $\delta = 0.05$ for individual kernel evaluations. The higher dimensionality supports reliable multi-scale testing across 7 bandwidth scales and 3 test types.

2. **Multi-scale bandwidths**: Features are distributed across $L$ bandwidth scales via divmod($D$, $L$), ensuring exact allocation with no feature loss and $O(D)$ total complexity.

3. **Numerical stability**: The computation $v_k = \sum_i y_i \cos(\cdot)$ is performed in float64 and remains stable in practice; in FlashS, centering and variance normalization are applied within the test channels used for inference.

### 3 Supplementary Note 3: Corrected Null Distribution Derivation

Under the null hypothesis of no spatial variation, the test statistic $T = \sum_{k=1}^{D} v_k^2$ where $v_k = \tilde{\mathbf{y}}^\top \mathbf{z}_k$ and $\tilde{\mathbf{y}}$ is the centered expression vector. Conditional on the spatial coordinates (and hence on $\mathbf{Z}$), the $v_k$ are linear combinations of $\tilde{\mathbf{y}}$ and are jointly normal under a Gaussian null.

#### 3.1 Covariance of the projection vector

The covariance matrix of the projection vector $\mathbf{v} = (v_1, \ldots, v_D)^\top$ is:

$$\mathbf{\Sigma}_v = \sigma_y^2 \, \mathbf{Z}_c^\top \mathbf{Z}_c / n \tag{20}$$

where $\mathbf{Z}_c$ denotes the column-centered feature matrix and $\sigma_y^2 = \mathrm{Var}[y_i]$ under the null.

#### 3.2 Why the standard Satterthwaite approximation fails

The standard approach assumes the $v_k$ are independent, giving $\mathrm{Var}[T] = 2 \sum_k \sigma_{v_k}^4$. However, at large bandwidth scales, the cosine features $z_k(\mathbf{s}) = \sqrt{2/D} \cos(\boldsymbol{\omega}_k^\top \mathbf{s} + b_k)$ with small $\|\boldsymbol{\omega}_k\|$ approximate similar low-frequency polynomials, creating non-negligible correlations. Accounting for these correlations under a Gaussian assumption on $\tilde{\mathbf{y}}$, the variance becomes:

$$\mathrm{Var}[T]\big|_{\mathrm{Gaussian}} = 2\|\mathbf{\Sigma}_v\|_F^2 = 2\sigma_y^4 \frac{1}{n^2} \|\mathbf{Z}_c^\top \mathbf{Z}_c\|_F^2 \tag{21}$$

This corrects for RFF feature correlations but still assumes $\tilde{y}_i$ is Gaussian; the additional correction for non-Gaussian (zero-inflated) expression is derived below.

#### 3.3 Per-scale decomposition

Features from different bandwidth scales are drawn from different spectral densities. At fine scales (small $\sigma$), cross-scale correlations are near-zero (mean $|r| < 0.01$), while at coarse scales (large $\sigma$), adjacent-scale correlations can be substantial (mean $|r| \approx 0.3$–$0.4$; Supplementary Fig. 5a). FlashS decomposes the Frobenius norm by scale:

$$\|\mathbf{Z}_c^\top \mathbf{Z}_c\|_F^2 \approx \sum_{\ell=1}^{L} \|\mathbf{Z}_{c,\ell}^\top \mathbf{Z}_{c,\ell}\|_F^2 \tag{22}$$

where $\mathbf{Z}_{c,\ell}$ contains the centered features for scale $\ell$. Crucially, this block-diagonal approximation does not affect the primary inference in FlashS: the multi-kernel Cauchy combination performs per-scale hypothesis tests, each using only within-scale features whose covariance $\mathbf{\Sigma}_{z,\ell} = \mathbf{Z}_{c,\ell}^\top \mathbf{Z}_{c,\ell}/n$ is computed exactly via the full per-scale Frobenius norm $\|\mathbf{\Sigma}_{z,\ell}\|_F^2$. This correction is substantial at large bandwidths—up to $\sim$15$\times$ the naive diagonal estimate $\sum_k \mathrm{Var}[z_k]^2$ (Supplementary Fig. 5b)—and is essential for correct null calibration. Under the Gaussian assumption, the per-scale decomposition yields:

$$\mathrm{Var}[T]\big|_{\mathrm{Gaussian}} \approx 2\sigma_y^4 n^2 \sum_{\ell=1}^{L} \|\mathbf{\Sigma}_{z,\ell}\|_F^2 \tag{23}$$

where $\mathbf{\Sigma}_{z,\ell} = \mathbf{Z}_{c,\ell}^\top \mathbf{Z}_{c,\ell}/n$ is the within-scale feature covariance.

##### 3.4 Kurtosis correction for non-Gaussian expression

The variance formula above holds under the implicit assumption that $\tilde{y}_i$ is Gaussian, so that the fourth central moment satisfies $\mathbb{E}[\tilde{y}^4] = 3\sigma_y^4$. In spatial transcriptomics, however, expression vectors are zero-inflated: the binary channel is Bernoulli, the rank channel is a mixture of zeros and ranks, and the direct channel inherits the zero-inflated count distribution. These distributions have excess kurtosis $\kappa_4 = \mathbb{E}[(\tilde{y}/\sigma_y)^4] - 3 \neq 0$.

For a quadratic form $T = \tilde{\mathbf{y}}^\top \mathbf{M} \tilde{\mathbf{y}}$ with independent $\tilde{y}_i$ having common variance $\sigma_y^2$ and excess kurtosis $\kappa_4$, the exact variance is[8]:

$$\mathrm{Var}[T] = 2\sigma_y^4 \|\mathbf{M}\|_F^2 + \kappa_4 \sigma_y^4 \sum_{i=1}^{n} M_{ii}^2 \tag{24}$$

where $\mathbf{M} = \mathbf{Z}_c \mathbf{Z}_c^\top$ and $M_{ii} = \|\mathbf{z}_{c,i}\|^2$ is the squared norm of the $i$-th row of the centered feature matrix. The second term vanishes under Gaussianity ($\kappa_4 = 0$) and is positive for leptokurtic distributions ($\kappa_4 > 0$), causing the Gaussian-only formula to underestimate $\mathrm{Var}[T]$ and hence produce p-values that are too small, inflating the false positive rate.

Applying the per-scale decomposition, the corrected per-scale variance is:

$$\mathrm{Var}[T_\ell] = 2\sigma_y^4 n^2 \|\mathbf{\Sigma}_{z,\ell}\|_F^2 + \kappa_4 \sigma_y^4 \sum_{i=1}^{n} \|\mathbf{z}_{c,\ell,i}\|^4 \tag{25}$$

where $\mathbf{z}_{c,\ell,i}$ is the centered feature vector of cell $i$ at scale $\ell$.

The per-gene excess kurtosis is computed analytically for the binary channel and numerically for the rank and direct channels:

- *Binary* ($y_i \in \{0,1\}$, $p = \mathrm{nnz}/n$): $\kappa_4 = (1 - 6p + 6p^2)/[p(1-p)]$.
- *Rank/Direct* (zero-inflated): $\kappa_4 = \frac{1}{n}\left[(n - \mathrm{nnz})\left(\frac{\mu}{\sigma}\right)^4 + \sum_{j\in\mathrm{nz}}\left(\frac{y_j-\mu}{\sigma}\right)^4\right] - 3$,

where $\mu$ and $\sigma$ are the population mean and standard deviation of the (zero-inclusive) expression vector.

At natural sparsity (∼90% zeros, typical of spatial transcriptomics), the binary channel has $\kappa_4 \approx$ 5–10 and the rank channel has $\kappa_4 \approx$ 15–25. Without the kurtosis correction, the missing term causes $\mathrm{Var}[T]$ to be underestimated by 10–30%, inflating FPR from the nominal 5% to 6–7%. Including the correction brings the simulated FPR to ∼5.5% at 90% sparsity, decreasing to ∼4.8% at 95% sparsity where the kurtosis term dominates.

##### 3.5 Moment-matching approximation

With $\mathbb{E}[T_\ell] = n\sigma_y^2 \sum_k \mathrm{Var}[z_{k,\ell}]$ and the kurtosis-corrected variance (Equation 25), we fit a scaled chi-square $\kappa\chi_\nu^2$ via:

$$\kappa = \frac{\mathrm{Var}[T_\ell]}{2\,\mathbb{E}[T_\ell]}, \qquad \nu = \frac{2\,\mathbb{E}[T_\ell]^2}{\mathrm{Var}[T_\ell]} \tag{26}$$

The p-value is then $p = \Pr[\kappa\chi_\nu^2 \geq T_{\mathrm{obs}}]$, computed via the survival function of the chi-square distribution.

##### 3.6 Efficient estimation

The within-scale covariance Frobenius norms $\|\mathbf{\Sigma}_{z,\ell}\|_F^2$ and the mean fourth power of centered row norms $\mathbb{E}[\|\mathbf{z}_{c,\ell,i}\|^4]$ are both estimated once on a coordinate subsample of size $M$ (default $M = 10{,}000$) in a single pass, at cost $O(M \sum_\ell D_\ell^2)$. This adds negligible overhead to the total computation and needs to be done only once per dataset. The population sum $\sum_i \|\mathbf{z}_{c,\ell,i}\|^4$ is estimated as $n \cdot \mathbb{E}[\|\mathbf{z}_{c,\ell,i}\|^4]$.

### 4 Supplementary Tables

| **Method** | **Mean $\tau$** | **Median $\tau$** | **Min $\tau$** | **$n$ datasets** |
|---|---|---|---|---|
| FlashS | **0.935** | **0.968** | 0.788 | 50 |
| SPARK-X[15] | 0.886 | 0.904 | 0.698 | 50 |
| nnSVG[13] | 0.798 | 0.815 | 0.301 | 49 |
| Moran's I | 0.777 | 0.800 | 0.207 | 50 |
| Spanve[2] | 0.775 | 0.869 | −0.365 | 50 |
| PreTSA$^{\dagger}$[16] | 0.769 | 0.803 | 0.261 | 50 |
| GPcounts[1] | 0.740 | 0.786 | 0.087 | 47 |
| SpaGFT[3] | 0.695 | 0.737 | 0.219 | 50 |
| SpatialDE2[6] | 0.678 | 0.738 | 0.059 | 49 |
| Hotspot$^{\dagger}$[4] | 0.654 | 0.699 | 0.202 | 50 |
| scBSP$^{\dagger}$[7] | 0.638 | 0.669 | −0.026 | 50 |
| SpatialDE[12] | 0.615 | 0.632 | 0.051 | 50 |
| SOMDE[5] | 0.571 | 0.609 | 0.082 | 44 |
| scGCO[14] | 0.514 | 0.541 | −0.062 | 50 |
| SPARK[11] | 0.420 | 0.494 | −0.455 | 42 |

Table 1: **Supplementary Table 1: SVG detection accuracy on the Open Problems benchmark.** Performance measured by Kendall $\tau$ correlation between predicted and ground truth spatial variability scores across 50 datasets spanning 9 platforms. Methods that failed on some datasets ($n < 50$) are scored only on datasets where they produced valid output. $^{\dagger}$PreTSA, Hotspot, and scBSP were not included in the original Open Problems leaderboard; we ran these methods on the same 50 datasets using their default parameters. SPARK-X's leaderboard value ($\tau = 0.886$) was independently validated by running SPARK-X in the same computing environment as FlashS ($\tau = 0.881$; Supplementary Table 10).

| Number of cells | Runtime (s) | Peak memory (GB) | Replicates |
|---|---|---|---|
| 1,000 | $4.7 \pm 1.6$ | $0.26 \pm 0.02$ | 3 |
| 5,000 | $7.2 \pm 1.4$ | $0.55 \pm 0.00$ | 3 |
| 10,000 | $9.7 \pm 0.2$ | $0.88 \pm 0.00$ | 3 |
| 25,000 | $19.8 \pm 0.0$ | $1.79 \pm 0.00$ | 3 |
| 50,000 | $37.7 \pm 0.1$ | $3.31 \pm 0.00$ | 3 |
| 100,000 | $74.0 \pm 0.5$ | $3.47 \pm 0.00$ | 3 |
| 200,000 | $145.4 \pm 1.6$ | $3.81 \pm 0.01$ | 3 |
| 500,000 | $366.2 \pm 13.3$ | $5.31 \pm 0.04$ | 3 |
| 1,000,000 | $703.8 \pm 3.9$ | $10.32 \pm 0.02$ | 3 |

Table 2: **Supplementary Table 2:** FlashS **scalability results.** Runtime and peak resident set size (RSS) for simulated datasets with 5,000 genes at varying cell counts. Values are mean ± s.d. across 3 independent replicates. Default parameters: $D = 500$ features, $L = 7$ scales. All experiments performed on a 16-core compute node.

### 5 Supplementary Figures

| Method | Runtime in seconds (mean ± s.d., $n = 3$) | | | | |
|---|---|---|---|---|---|
| | **1,000** | **5,000** | **10,000** | **25,000** | **50,000** |
| PreTSA[16] | $0.1 \pm 0.0$ | $0.3 \pm 0.1$ | $0.5 \pm 0.0$ | $1.2 \pm 0.1$ | $2.4 \pm 0.1$ |
| scBSP[7] | $0.8 \pm 0.1$ | $2.4 \pm 0.1$ | $4.4 \pm 0.2$ | $10.6 \pm 0.4$ | $20.1 \pm 0.3$ |
| Moran's I (Squidpy[9]) | $5.6 \pm 8.6$ | $1.0 \pm 0.1$ | $2.6 \pm 0.9$ | $4.6 \pm 0.5$ | $8.3 \pm 2.0$ |
| FlashS | $6.7 \pm 3.9$ | $9.1 \pm 2.1$ | $17.2 \pm 1.3$ | $37.4 \pm 3.9$ | $67.1 \pm 2.9$ |
| SPARK-X[15] | $9.4 \pm 0.4$ | $10.5 \pm 0.2$ | $11.5 \pm 0.2$ | $15.3 \pm 0.7$ | $19.9 \pm 0.9$ |
| Spanve[2] | $9.5 \pm 0.1$ | $20.8 \pm 0.3$ | $20.8 \pm 0.6$ | $29.9 \pm 0.6$ | $39.7 \pm 0.3$ |
| Hotspot[4] | $30.1 \pm 1.5$ | $34.4 \pm 1.6$ | $45.4 \pm 0.7$ | $68.0 \pm 9.1$ | $108.8 \pm 5.1$ |
| SOMDE[5] | $97.9 \pm 0.2$ | $145.6 \pm 3.0$ | $167.6 \pm 4.0$ | $145.0 \pm 2.2$ | $152.2 \pm 0.5$ |
| SpatialDE[12] | $121.2 \pm 3.3$ | $658.7 \pm 24.9$ | $3{,}777 \pm 103$ | timeout | OOM |
| nnSVG[13] | $6{,}603 \pm 185$ | timeout | timeout | timeout | timeout |

| Method | Peak memory in MB (mean, $n = 3$) | | | | |
|---|---|---|---|---|---|
| | **1,000** | **5,000** | **10,000** | **25,000** | **50,000** |
| PreTSA | 358 | 1,105 | 1,983 | 4,628 | 9,031 |
| scBSP | 679 | 1,388 | 2,277 | 4,883 | 9,257 |
| Moran's I | 599 | 807 | 929 | 1,883 | 3,469 |
| FlashS | 570 | 807 | 929 | 1,882 | 3,470 |
| SPARK-X | 781 | 1,791 | 3,077 | 6,942 | 13,379 |
| Spanve | 601 | 980 | 1,123 | 2,075 | 3,662 |
| Hotspot | 747 | 806 | 929 | 1,883 | 3,470 |
| SOMDE | 594 | 805 | 1,123 | 2,075 | 3,662 |
| SpatialDE | 682 | 3,414 | 11,515 | timeout | OOM |
| nnSVG | 1,108 | timeout | timeout | timeout | timeout |

Table 3: **Supplementary Table 3: Runtime and memory comparison across SVG detection methods.** Simulated datasets with 5,000 genes at varying cell counts. Runtime values are mean ± s.d. across 3 replicates; peak memory is mean peak resident set size. Each method–size–replicate combination was run as an isolated subprocess with a 2-hour timeout and 50 GB memory limit. Methods are sorted by runtime at 5,000 cells. "timeout" indicates the method exceeded the 2-hour limit; "OOM" indicates out-of-memory termination. nnSVG required ∼6,600 s (∼110 min) at 1,000 cells with 5,000 genes, and exceeded the 2-hour timeout at all larger cell counts (5,000–50,000), as the nearest-neighbor Gaussian process model fits each gene individually with iterative REML optimization. FlashS runtimes in this table are slightly higher than in Supplementary Table 2 because each method–size–replicate combination was launched as an independent subprocess with startup overhead. All experiments performed on a 16-core compute node (AMD EPYC).

| | FlashS | | | PreTSA | | |
|---|---|---|---|---|---|---|
| **Cells** | **Time (s)** | **Mem (GB)** | $\tau$ | **Time (s)** | **Mem (GB)** | $\tau$ |
| 1,000 | $4.7 \pm 1.6$ | 0.3 | | $0.1 \pm 0.0$ | 0.4 | |
| 5,000 | $7.2 \pm 1.4$ | 0.6 | | $0.3 \pm 0.1$ | 1.1 | |
| 10,000 | $9.7 \pm 0.2$ | 0.9 | | $0.5 \pm 0.0$ | 2.0 | |
| 25,000 | $19.8 \pm 0.0$ | 1.8 | | $1.2 \pm 0.1$ | 4.6 | |
| 50,000 | $37.7 \pm 0.1$ | 3.3 | **0.935** | $2.4 \pm 0.1$ | 9.0 | 0.769 |
| 100,000 | $74.0 \pm 0.5$ | 3.5 | | $4.6 \pm 0.0$ | 17.8 | |
| 200,000 | $145.4 \pm 1.6$ | 3.8 | | $8.6 \pm 0.2$ | 27.6 | |
| 500,000 | $366.2 \pm 13.3$ | 5.3 | | $21.5 \pm 0.4$ | 68.7 | |
| 1,000,000 | $703.8 \pm 3.9$ | 10.3 | | $43.7 \pm 0.6$ | 137.0 | |

Table 4: **Supplementary Table 4: Extended scalability comparison between** FlashS **and PreTSA.** Runtime (mean ± s.d., $n = 3$) and peak memory for simulated datasets with 5,000 genes across cell counts from 1,000 to 1,000,000. PreTSA is faster in wall-clock time at all scales due to optimized dense BLAS operations on the shared hat matrix, but its memory scales as $O(n \cdot g)$ with the dense expression matrix: 137 GB at 1,000,000 cells compared to 10.3 GB for FlashS. On workstations with ≤64 GB RAM, PreTSA becomes infeasible beyond ∼200,000 cells. Detection accuracy ($\tau$) on the Open Problems benchmark (50 datasets) is shown for reference.

| **Category** | **Genes** | **GO terms** | **Top pathway** | **Top $p$-value** |
|---|---|---|---|---|
| Both (FlashS ∩ PreTSA) | 1,030 | 440 | Oxidative phosphorylation | $3.91 \times 10^{-72}$ |
| FlashS-only | 2,669 | 305 | Mitochondrial translation elongation | $6.71 \times 10^{-14}$ |
| PreTSA-only | 93 | 0 | — | — |
| Neither | 10,842 | — | — | — |

Table 5: **Supplementary Table 5: SVG detection categories and GO enrichment summary on Visium human heart.** Gene counts by detection category among 14,634 genes tested by both methods, using Benjamini–Hochberg $q < 0.05$ separately within method. Gene Ontology enrichment used the one-sided g:Profiler over-representation test with the 14,634-gene universe as the custom background and g:SCS correction across terms. Exact corrected p-values for the pathways shown are $3.91 \times 10^{-72}$ for oxidative phosphorylation among shared SVGs and $6.71 \times 10^{-14}$ for mitochondrial translation elongation among FlashS-only SVGs; all exact term-level p-values are provided in Supplementary Data 1. PreTSA-only SVGs yielded no significant term after correction.

| **Configuration** | **Normalize** | **Log** | **Mean $\tau$** | **Median $\tau$** | **Min $\tau$** | $\Delta\tau$ |
|---|---|---|---|---|---|---|
| Raw counts (default) | No | No | **0.935** | **0.968** | 0.788 | — |
| Log-transform only | No | Yes | 0.935 | 0.968 | 0.774 | −0.001 |
| Normalization only | Yes | No | 0.779 | 0.814 | 0.133 | −0.157 |
| Norm. + log | Yes | Yes | 0.787 | 0.818 | 0.150 | −0.148 |

Table 6: **Supplementary Table 6: Preprocessing ablation results.** Detection accuracy across 50 Open Problems datasets for four preprocessing configurations. Library-size normalization is the primary factor reducing accuracy ($\Delta\tau \approx -0.15$), while log-transformation has negligible impact ($\Delta\tau = -0.001$). All configurations use identical model parameters ($D = 500$, $L = 7$).

| Gene set | SVG category | Size | Overlap | Expected | Fold | OR (95% CI) | $Q$ (BH) | Sig. |
|---|---|---|---|---|---|---|---|---|
| PGC-1$\alpha$ targets (78) | FlashS-only (2,669) | 78 | 27 | 14.2 | 1.90 | 2.39 (1.51–3.83) | $1.0 \times 10^{-3}$ | ✓ |
| | Both (1,030) | 78 | 15 | 5.5 | 2.73 | 3.18 (1.86–5.69) | $1.0 \times 10^{-3}$ | ✓ |
| | PreTSA-only (93) | 78 | 0 | 0.5 | 0 | — | 1.00 | |
| | Neither (10,842) | 78 | 36 | 57.8 | 0.62 | 0.35 (0.23–0.53) | 1.00 | |
| OXPHOS hallmark (199) | FlashS-only | 199 | 56 | 36.3 | 1.54 | 1.77 (1.31–2.43) | $1.0 \times 10^{-3}$ | ✓ |
| | Both | 199 | 109 | 14.0 | 7.78 | 17.8 (13.3–23.6) | $1.0 \times 10^{-71}$ | ✓ |
| | PreTSA-only | 199 | 0 | 1.3 | 0 | — | 1.00 | |
| | Neither | 199 | 34 | 147.4 | 0.23 | 0.08 (0.06–0.12) | 1.00 | |
| Mito biogenesis (49) | FlashS-only | 49 | 39 | 8.9 | 4.36 | 17.7 (8.65–33.8) | $9.5 \times 10^{-20}$ | ✓ |
| | Both | 49 | 1 | 3.5 | 0.29 | 0.27 (0.07–2.20) | 1.00 | |
| | PreTSA-only | 49 | 0 | 0.3 | 0 | — | 1.00 | |
| | Neither | 49 | 9 | 36.3 | 0.25 | 0.12 (0.06–0.27) | 1.00 | |
| DCM KEGG (71) | FlashS-only | 71 | 13 | 12.9 | 1.00 | 1.01 (0.56–1.87) | 1.00 | |
| | Both | 71 | 28 | 5.0 | 5.60 | 8.81 (5.50–14.3) | $4.6 \times 10^{-14}$ | ✓ |
| | PreTSA-only | 71 | 0 | 0.5 | 0 | — | 1.00 | |
| | Neither | 71 | 30 | 52.6 | 0.57 | 0.31 (0.20–0.50) | 1.00 | |
| Myogenesis hallmark (177) | FlashS-only | 177 | 58 | 32.3 | 1.80 | 2.21 (1.62–3.05) | $8.8 \times 10^{-6}$ | ✓ |
| | Both | 177 | 66 | 12.5 | 5.30 | 8.32 (6.12–11.4) | $2.4 \times 10^{-30}$ | ✓ |
| | PreTSA-only | 177 | 0 | 1.1 | 0 | — | 1.00 | |
| | Neither | 177 | 53 | 131.1 | 0.40 | 0.19 (0.14–0.27) | 1.00 | |

Table 7: **Supplementary Table 7: Heart metabolic pathway enrichment analysis with Benjamini–Hochberg correction.** Hypergeometric enrichment of SVG categories against five curated gene sets from MSigDB. All methods use unified Benjamini–Hochberg correction at $q < 0.05$. "Size" is the gene set size in the universe of 14,634 tested genes. "Fold" is fold enrichment (observed/expected). OR is the odds ratio with 95% confidence interval from Fisher's exact test. $Q$ (BH) is the Benjamini–Hochberg adjusted p-value across all 20 tests (5 gene sets × 4 SVG categories). ✓ indicates significance at FDR < 0.05. Category sizes are shown in parentheses. Of 49 curated mitochondrial biogenesis genes, 39 are detected exclusively by FlashS. PreTSA-only SVGs (93 genes) show zero overlap with every tested gene set (0 of 20 tests significant).

| Sample | Common | Both | FlashS-unique | SPARK-X-unique | FlashS GO | Top term (FlashS-unique) | SPARK-X GO |
|---|---|---|---|---|---|---|---|
| *Donor 1* | | | | | | | |
| 151507 | 13,329 | 4,697 | 5,043 | 43 | 7 | chromatin organization ($3.7 \times 10^{-4}$) | 0 |
| 151508 | 12,863 | 4,326 | 4,297 | 47 | 1 | Pre-mRNA processing ($4.8 \times 10^{-2}$) | 0 |
| 151509 | 13,693 | 3,737 | 6,641 | 26 | 24 | nucleoplasm ($1.2 \times 10^{-8}$) | 0 |
| 151510 | 13,398 | 4,840 | 4,949 | 43 | 4 | chromatin remodeling ($8.1 \times 10^{-4}$) | 0 |
| *Donor 2* | | | | | | | |
| 151669 | 13,600 | 9,716 | 2,314 | 40 | 18 | DNA-binding TF activity ($7.0 \times 10^{-14}$) | 0 |
| 151670 | 13,251 | 9,013 | 2,438 | 31 | 15 | DNA-binding TF activity ($9.1 \times 10^{-11}$) | 0 |
| 151671 | 14,032 | 10,298 | 1,366 | 291 | 13 | DNA-binding TF activity ($4.9 \times 10^{-5}$) | 0 |
| 151672 | 13,711 | 8,592 | 2,040 | 233 | 0 | — | 0 |
| *Donor 3* | | | | | | | |
| 151673 | 14,248 | 11,492 | 1,248 | 50 | 6 | DNA-binding TF activity ($9.2 \times 10^{-5}$) | 1 |
| 151674 | 15,094 | 12,296 | 1,085 | 79 | 11 | DNA-binding TF activity, RNAPII ($2.3 \times 10^{-7}$) | 0 |
| 151675 | 13,727 | 11,418 | 1,002 | 53 | 13 | DNA-binding TF activity ($3.0 \times 10^{-4}$) | 0 |
| 151676 | 13,797 | 10,982 | 1,204 | 70 | 15 | DNA-binding TF activity ($2.1 \times 10^{-10}$) | 0 |
| Mean | 13,812 | 8,451 | 2,802 | 84 | 10.6 | | 0.1 |

Table 8: **Supplementary Table 8: DLPFC per-sample GO enrichment of method-specific SVGs.** For each of the 12 DLPFC Visium samples, genes tested by both FlashS and SPARK-X form the common gene universe. SPARK-X p-values are corrected via Benjamini–Hochberg; genes are categorized at $q < 0.05$. GO enrichment of method-specific gene sets was performed via g:Profiler with the per-sample common gene universe as custom background (GO:BP, GO:MF, GO:CC, KEGG, Reactome; g:SCS threshold $p < 0.05$). "FlashS GO" and "SPARK-X GO" denote the number of significant enrichment terms. FlashS-unique SVGs are enriched for DNA-binding transcription factor activity in 7 of 12 samples ($p < 10^{-4}$–$10^{-14}$), while SPARK-X-unique SVGs yield at most 1 marginally enriched term (sample 151673, C-type lectin signaling). The asymmetry is consistent across all three donors.

| Config | Permutation scheme | Preserved structure | Replicates | Rejection rate |
|---|---|---|---|---|
| N0 | Global row permutation | None | 3 | 5.6% ± 1.1% |
| N1 | Within-section permutation | Between-section | 10 | 100% |
| N2 | Spatial block ($6 \times 6$ grid) | Within-block | 6 | 100% |

Table 9: **Supplementary Table 9: Structured null testing on the Allen MERFISH atlas.** Rejection rate (fraction of 550 genes with $p < 0.05$) under three permutation strategies applied to 3,938,808 cells. N0 destroys all spatial structure and serves as a calibration control; the observed 5.6% FPR is close to the nominal 5% level (Fig. 3g–i). N1 retains between-section composition (59 brain sections with distinct expression profiles); N2 retains within-block spatial autocorrelation at scale ∼1/6 of each section diameter. Under both structured schemes, 100% rejection is expected because genuine spatial patterns persist at preserved scales. The contrast between N0 (5.6%) and N1/N2 (100%) supports the interpretation that FlashS's detections arise from real spatial organization rather than method miscalibration.

| Method | Mean $\tau$ | Median $\tau$ | Datasets | Win rate |
|---|---|---|---|---|
| FlashS | **0.935** | **0.968** | 50 | **50/50** |
| SPARK-X | 0.881 | 0.900 | 50 | 0/50 |

Table 10: **Supplementary Table 10: Same-environment SVG ranking comparison between FlashS and SPARK-X on the Open Problems benchmark.** Both methods were run on all 50 datasets in the same computing environment (identical data loading, evaluation pipeline, and hardware). SPARK-X was run via the R package SPARK (v1.1.1) using default parameters (`option = "mixture"`). FlashS was run with default parameters ($D = 500$, $L = 7$, three-part test on raw counts). The SPARK-X result ($\tau = 0.881$) is consistent with the Open Problems leaderboard value ($\tau = 0.886$), validating the fairness of the comparison. FlashS outperforms SPARK-X on every dataset, with the largest per-dataset advantages on tissues with complex multi-scale spatial architecture (cerebellum $\Delta\tau = +0.113$, skin melanoma +0.113, heart +0.099).

| Method | Kernel type | Detected | Fraction | Genome-wide SVGs |
|---|---|---|---|---|
| FlashS | Multi-scale Gaussian (RFF) | 40/49 | 82% | 3,699/14,634 (25%) |
| Moran's I | Single-scale autocorrelation | 28/49 | 57% | 18,850/36,601 (51%) |
| SPARK-X | Periodic covariance projection | 25/49 | 51% | 2,264/14,634 (15%) |
| PreTSA | B-spline tensor product | 1/49 | 2% | 1,123/14,634 (8%) |

Table 11: **Supplementary Table 11: Detection of 49 mitochondrial biogenesis genes across SVG methods on Visium human heart.** "Detected" counts genes significant at $q < 0.05$ (Benjamini–Hochberg correction for all methods) among the 49 curated mitochondrial biogenesis genes (composition: 21 large-subunit mitoribosomal proteins [*MRPL1, 2, 4, 11, 13, 14, 16, 17, 20, 22, 24, 28, 35, 37, 40, 47, 51, 53, 54, 57, 58*]; 12 small-subunit mitoribosomal proteins [*MRPS5, 6, 7, 11, 15, 17, 18B, 24, 25, 26, 34, 35*]; 7 TOM/TIM translocase components [*TOMM20, 22, 34, 40L, TIMM17B, 21, 44*]; 7 respiratory chain assembly factors [*NDUFAF1, AF2, AF4, NDUFA5, A9, A10, NDUFB5*]; and 2 cytochrome c oxidase assembly components [*COX10, COX11*]). "Genome-wide SVGs" indicates the total number of significant genes detected by each method. FlashS, SPARK-X, and PreTSA were tested on 14,634 commonly expressed genes; Moran's I was computed via Squidpy ($k = 6$ nearest neighbors) on all 36,601 genes in the expression matrix (including unexpressed genes that receive valid p-values under Squidpy 1.6.1). SPARK-X detects 51% of the module from a 15% genome-wide call rate, Moran's I detects 57% from a 51% call rate, and FlashS detects 82% from a 25% call rate. The detection gradient is consistent with multi-scale kernel methods recovering more of this compound-pattern gene set.

| Method | Hallmark ($n = 50$) Mean | Hallmark ($n = 50$) Median | KEGG ($n = 315$) Mean | KEGG ($n = 315$) Median | Overall ($n = 365$) Mean | Overall ($n = 365$) Median |
|---|---|---|---|---|---|---|
| FlashS | **0.405** | **0.392** | **0.357** | **0.338** | **0.363** | **0.347** |
| Moran's I | 0.335 | 0.315 | 0.306 | 0.286 | 0.310 | 0.289 |
| PreTSA | 0.149 | 0.135 | 0.137 | 0.117 | 0.139 | 0.118 |
| SPARK-X | 0.266 | 0.247 | 0.229 | 0.201 | 0.234 | 0.209 |

Table 12: **Supplementary Table 12: Module completeness of SVG detection across 365 curated gene sets on Visium human heart.** For each gene set (50 MSigDB Hallmark and 315 KEGG pathways with $\geq$5 genes in the 14,634-gene universe), module completeness is defined as the fraction of pathway genes detected as SVGs at $q < 0.05$ (Benjamini–Hochberg, applied uniformly to all methods). All four methods are compared within the same 14,634-gene universe; for Moran's I, genome-wide BH-corrected SVGs are intersected with this universe (3,006 SVGs). FlashS achieves the highest completeness across all pathway categories (mean 0.363 overall), surpassing even Moran's I (0.310) despite both methods detecting comparable numbers of SVGs within this universe (3,699 vs 3,006). This indicates that FlashS's multi-kernel approach selectively detects pathway-relevant spatial genes. PreTSA achieves the lowest completeness (0.149 on Hallmark), while SPARK-X (0.266 on Hallmark) falls between Moran's I and PreTSA, consistent across all 365 pathways (Supplementary Fig. 10).

Table 13: **Supplementary Table 13: Ranking-metric ablation on the Open Problems benchmark.** Mean Kendall $\tau$ across all 50 datasets for two ranking strategies applied to FlashS output. The spatial effect size (default) is the maximum ratio of observed to expected test statistic across the test channels; $-\log_{10}(p)$-only uses p-values alone. The default effect size yields substantially higher concordance ($\tau = 0.935$) than $-\log_{10}(p)$ alone ($\tau = 0.868$). This pattern reflects the well-known limitation of p-value-based rankings: at moderate-to-large sample sizes, p-values saturate near machine precision, collapsing discriminative power. The effect size provides a continuous, non-saturating ranking metric with a direct statistical interpretation (signal-to-noise ratio). Note that most leaderboard methods also use effect-size-type scores rather than p-values: SpatialDE, SpatialDE2, and SOMDE report fraction of spatial variance (FSV); Moran's I reports the I statistic; SpaGFT and Sepal report method-specific test statistics.

| Score | Mean $\tau$ | Median $\tau$ | $\Delta$ vs effect size |
|---|---|---|---|
| Effect size (default) | 0.935 | 0.968 | — |
| $-\log_{10}(p)$ only | 0.868 | 0.879 | −0.067 |

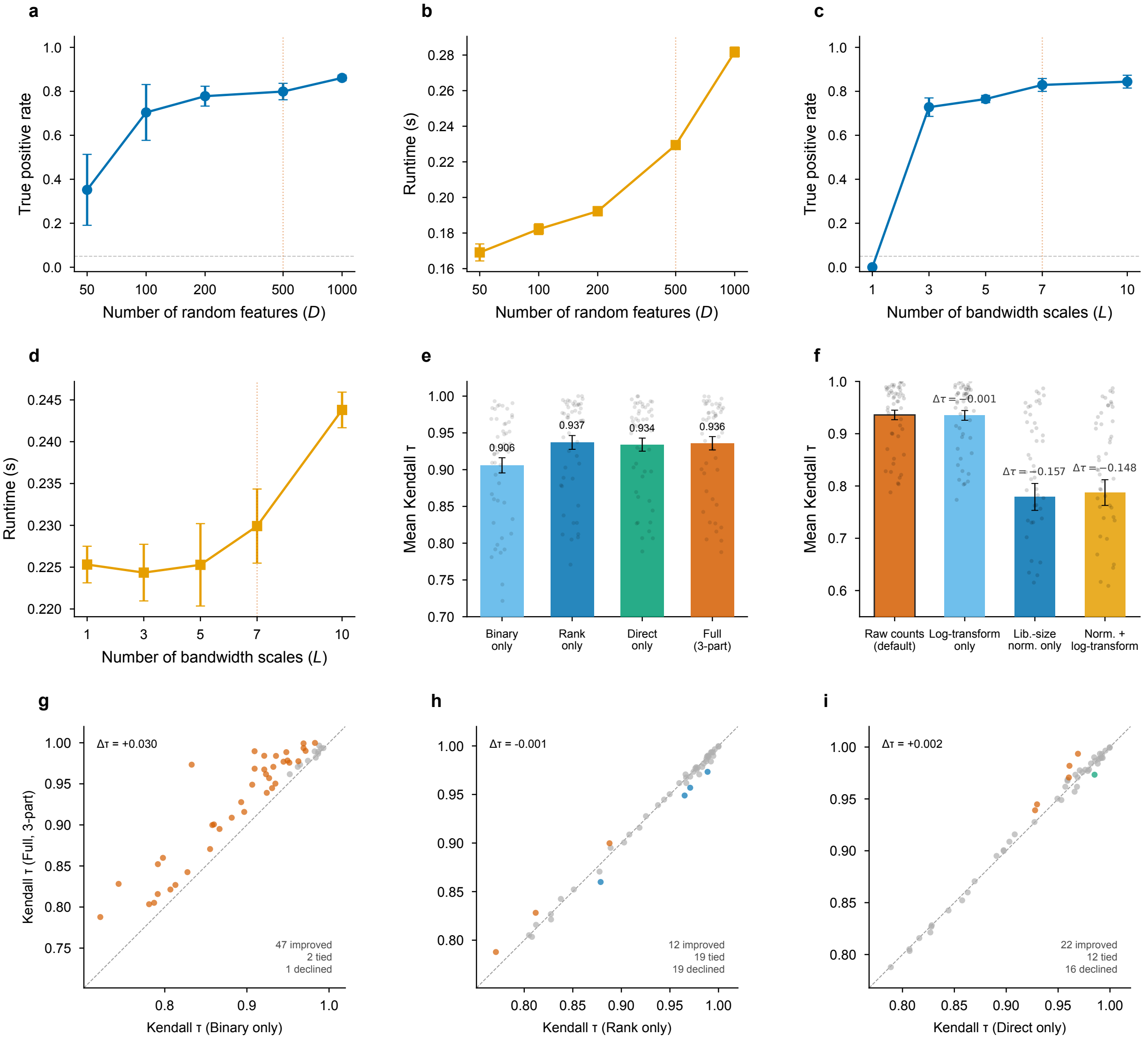

Figure 1: **Supplementary Figure 1: Method robustness—hyperparameter sensitivity and ablation studies. Row 1—Hyperparameter sensitivity: (a)** Detection power (true positive rate at $q < 0.05$) as a function of the number of random features $D$, with $L = 7$ bandwidth scales. **(b)** Runtime per 400 genes as a function of $D$. **(c)** Detection power as a function of the number of bandwidth scales $L$, with $D = 500$. Dotted red lines indicate default values ($D = 500$, $L = 7$). Simulations use 200 spatially variable genes (hotspot pattern, effect size 0.3, 30% sparsity) and 200 null genes across 2,500 cells; error bars indicate standard error over 3 replicates. **Row 2—Ablation and preprocessing: (d)** Runtime per 400 genes as a function of $L$. **(e)** Mean Kendall $\tau$ across 50 Open Problems datasets for each test configuration: binary-only, rank-only, direct-only, and full three-part test. Gray dots show individual datasets; error bars indicate SEM. **(f)** Effect of preprocessing on detection accuracy. $\Delta\tau$ relative to raw counts (default). **Row 3—Per-dataset paired comparisons: (g–i)** Paired Kendall $\tau$ between the full model and each single-channel configuration (binary, rank, direct) across all 50 datasets. Points above the diagonal indicate datasets where the full model outperforms the single channel. Counts of improved / tied / declined datasets are annotated.

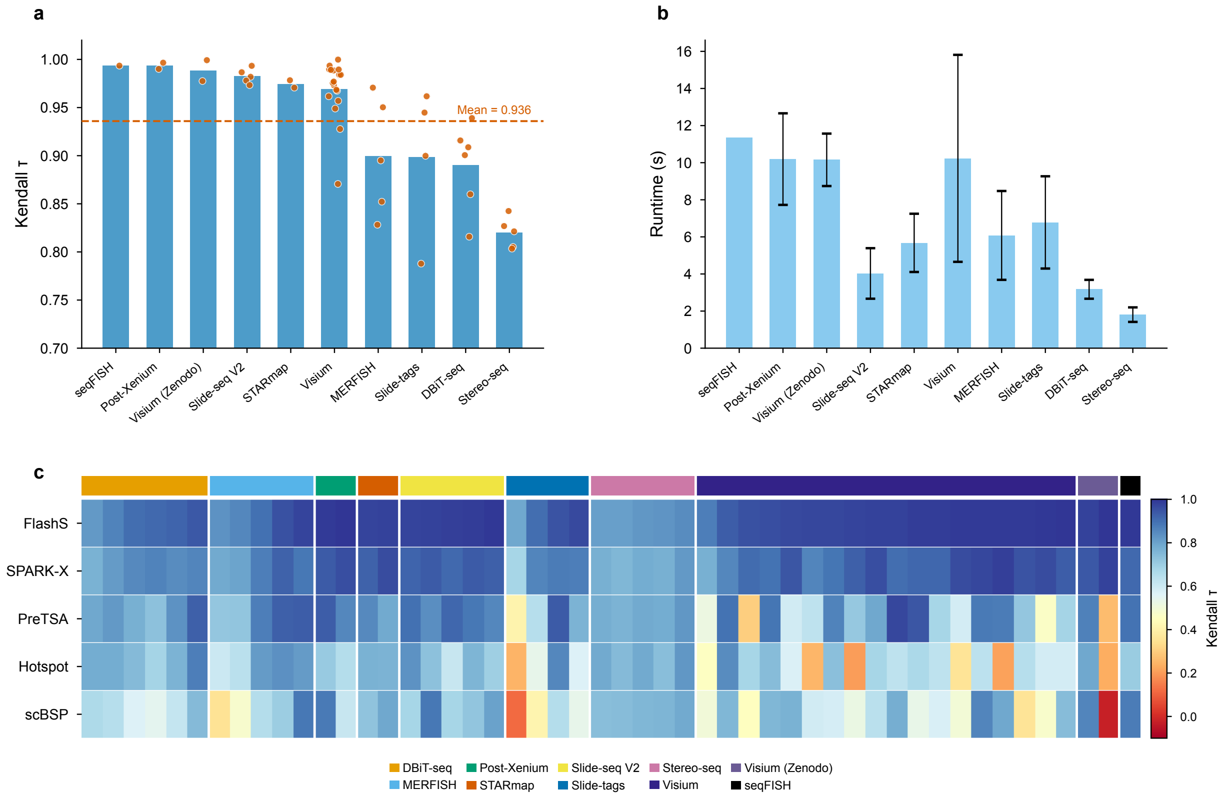

Figure 2: **Supplementary Figure 2: Benchmark performance heterogeneity across platforms and datasets. (a)** Mean Kendall $\tau$ correlation for each of the 9 spatial transcriptomics platforms, sorted by performance. Individual dataset values shown as dots; error bars indicate standard error. Dashed red line marks overall mean $\tau = 0.935$. **(b)** Mean runtime per dataset by platform, sorted by runtime. FlashS completes in under 25 seconds per dataset. **(c)** Kendall $\tau$ for FlashS, SPARK-X, PreTSA, Hotspot, and scBSP across all 50 Open Problems datasets, grouped by platform (colored bar) and sorted by FlashS performance within each group. FlashS achieves $\tau > 0.78$ on all datasets. SPARK-X (mean $\tau = 0.881$) ranks second but falls below FlashS on every dataset ($\Delta\tau = +0.055$). PreTSA shows intermediate accuracy (mean $\tau = 0.769$); Hotspot and scBSP show substantially lower and more variable accuracy, particularly on Slide-seq and STARmap platforms.

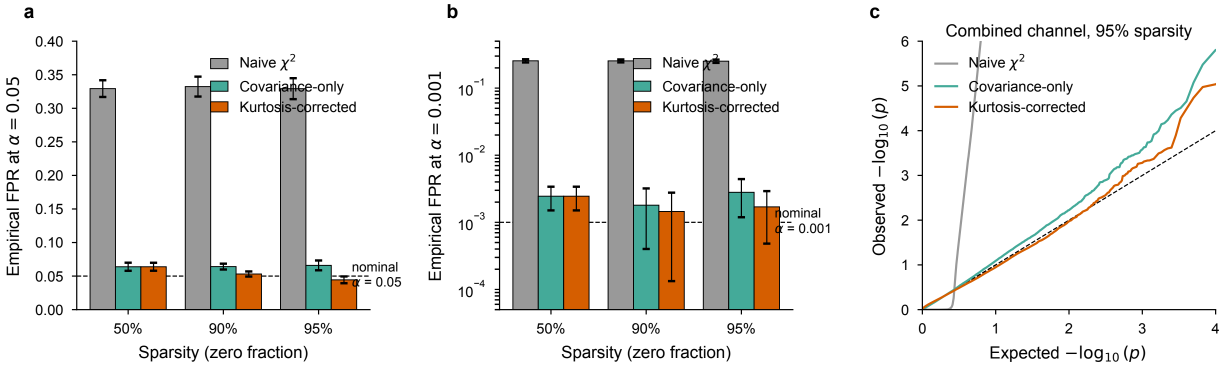

Figure 3: **Supplementary Figure 3: Kurtosis-correction ablation.** The same per-gene test statistics computed on null simulations ($50\times50$ grid; sparsities $\{50\%, 90\%, 95\%\}$; 20 replicates $\times$ 1,000 genes) are evaluated under three Satterthwaite null variance formulas: naive $\chi^2(D)$ assuming i.i.d. Gaussian RFF projections (grey); covariance-only Gaussian quadratic form $\mathrm{Var}[T] = 2\sigma^4 n^2 \|\mathrm{Cov}_s\|_F^2$ (teal); and kurtosis-corrected $\mathrm{Var}[T] = 2\sigma^4 n^2 \|\mathrm{Cov}_s\|_F^2 + \kappa_4 \sigma^4 n \sum_i \|\mathbf{z}_{c,s,i}\|^4$ (vermillion; FlashS default). **(a)** Empirical FPR at $\alpha = 0.05$, combined channel. The naive $\chi^2$ baseline is inflated by $\sim$6.6$\times$ across all sparsities, confirming that RFF column correlations cannot be ignored. Covariance correction brings FPR close to nominal; the kurtosis term then removes residual inflation that emerges at high sparsity ($0.066 \to 0.044$ at 95% zeros). **(b)** FPR at the extreme-tail threshold $\alpha = 10^{-3}$ (log scale). Here the kurtosis correction is most visible: covariance-only yields 2.8$\times$ inflation at 95% sparsity, while the kurtosis-corrected test stays within 1.7$\times$ of nominal. Naive $\chi^2$ fails by two orders of magnitude. **(c)** QQ plot of pooled null p-values at 95% sparsity, combined channel. The kurtosis-corrected curve tracks the diagonal closely into $-\log_{10}(p) = 4$; covariance-only shows mild upward deviation; naive $\chi^2$ diverges sharply. Error bars in (a,b) show standard deviation across 20 replicates.

| **Dataset** | $\tau$ | $\rho$ | **J@200** | **J@500** | **SVGs (raw)** | **SVGs (norm)** |
|---|---|---|---|---|---|---|
| Heart (4,235 spots) | 0.751 | 0.891 | 0.569 | 0.739 | 3,699 | 4,098 |
| DLPFC 151673 | 0.921 | 0.988 | 0.329 | — | 12,677 | 12,369 |
| DLPFC 151669 | 0.944 | 0.989 | 0.159 | — | 11,962 | 11,534 |

Table 14: **Supplementary Table 14: SVG ranking stability after library-size normalization.** FlashS was re-run with `normalize=True` (library-size normalization to median total counts per spot) on raw counts. Rankings are compared against the default FlashS configuration (`normalize=False`) using Kendall $\tau$ and Spearman $\rho$ on effect-size ranks, and Jaccard overlap at top-$k$. SVG counts use unified Benjamini–Hochberg correction across the 14,634-gene universe (heart) or per-sample tested genes (DLPFC), consistent with the main text. DLPFC rankings are highly concordant ($\tau > 0.92$), and cortical layer marker enrichment is preserved. Heart rankings show moderate concordance ($\tau = 0.75$), reflecting the fact that library size in cardiac tissue is partly spatially structured (cardiomyocyte-rich regions have higher total RNA content). Both DLPFC samples are from different donors to assess generalizability (151673: Donor 3; 151669: Donor 2).

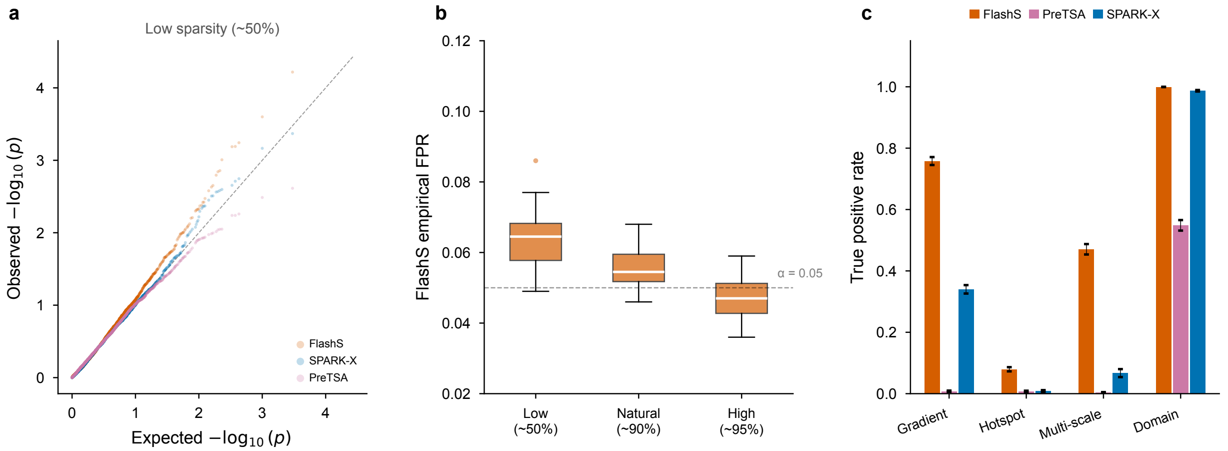

Figure 4: **Supplementary Figure 4: Extended calibration and detection robustness.** Cross-method FPR comparison and QQ plots at natural and high sparsity are shown in main text Fig. 3a–c; this figure provides complementary detail. **(a)** QQ plot of null p-values at low sparsity (∼50% zeros). All three methods (FlashS, SPARK-X, PreTSA) track the diagonal closely. **(b)** FlashS empirical FPR at $\alpha = 0.05$ across three zero-fraction levels (∼50%, ∼90%, ∼95%; 2,500 spots, 20 replicates each). With the kurtosis-corrected Satterthwaite approximation (Supplementary Note 3), mean FPR is 5.6%, close to the nominal level (dashed line) and improving at higher sparsity (4.8% at ∼95% zeros). **(c)** Detection power (true positive rate at $q < 0.05$) for FlashS, PreTSA, and SPARK-X across four spatial pattern types (gradient, hotspot, multiscale, domain) at effect size 1.0 and 80% sparsity. This complements Fig. 3d (50% sparsity) by showing how power degrades at higher zero fractions. FlashS retains the highest power on hotspot and multiscale patterns under high sparsity. Error bars indicate standard error across 10 replicates.

| $D$ | **Time (s)** | **Speedup** | **$\tau$ vs $D$=500** | **J@100** | **Blank Q4** | **Markers@100** |
|---|---|---|---|---|---|---|
| 500 | 755 | 1.0× | ref | ref | 82% | 4/12 |
| 200 | 559 | 1.4× | 0.958 | 0.905 | 80% | 4/12 |
| 100 | 340 | 2.2× | 0.882 | 0.739 | 86% | 4/12 |
| 50 | 222 | 3.4× | 0.900 | 0.786 | 82% | 5/12 |
| PreTSA | 25 | — | — | — | 90% | 5/12 |

Table 15: **Supplementary Table 15: Ranking stability at reduced feature counts on the Allen MERFISH atlas.** FlashS was run on the full MERFISH dataset (3,938,808 cells × 550 genes) at $D \in \{50, 100, 200\}$ with $L = 7$ fixed. "$\tau$ vs $D = 500$" is Kendall $\tau$ between effect-size rankings at reduced $D$ and the $D = 500$ baseline. "Blank bot. Q" is the percentage of 50 blank negative controls in the bottom quartile of the ranking. "Markers in top-100" counts known neurotransmitter/glial markers in the top 100. At $D = 50$ (222 s), FlashS retains $\tau = 0.900$ relative to the $D = 500$ baseline, with identical blank demotion (82%) and comparable marker retention. PreTSA completes in 25 s on this dataset; FlashS at $D = 50$ is slower but preserves the ranking fidelity of the full $D = 500$ model. At $D = 200$, $\tau = 0.958$ and Jaccard@100 = 0.905, indicating near-identical rankings. This confirms that reducing $D$ for faster runtime produces graceful degradation rather than qualitative ranking changes.

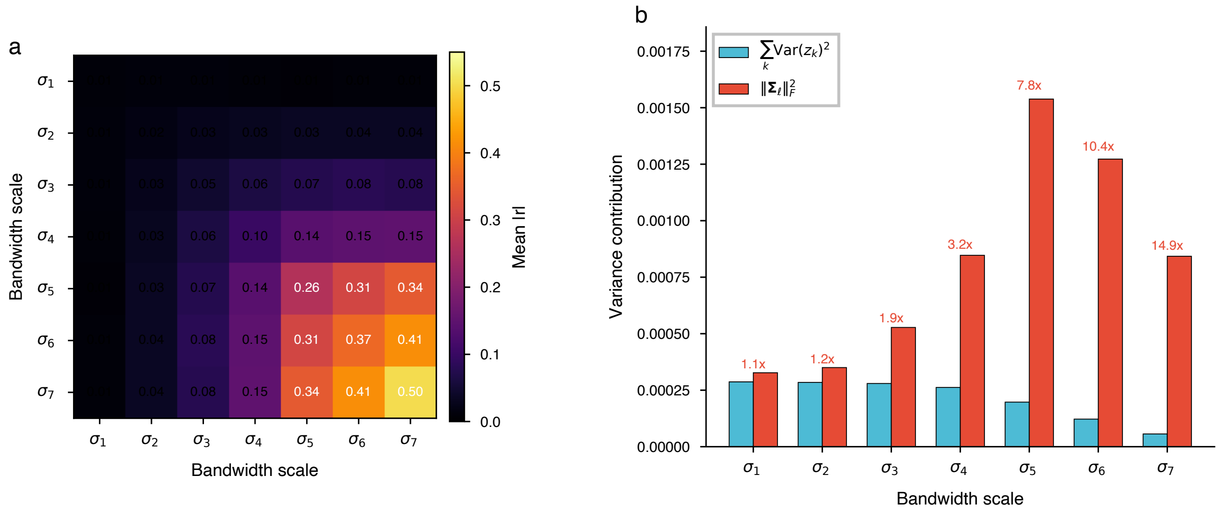

Figure 5: **Supplementary Figure 5: Cross-scale RFF feature correlation structure and within-scale covariance correction.** **(a)** Mean absolute Pearson correlation $|r|$ between RFF feature blocks at each pair of bandwidth scales ($L = 7$, $D = 500$) on a $50 \times 50$ regular grid. Fine-scale features ($\sigma_1$–$\sigma_3$) show near-zero cross-scale correlations (mean $|r| < 0.08$). Coarser scales ($\sigma_5$–$\sigma_7$) exhibit non-negligible cross-scale correlations (mean $|r| \approx 0.3$–$0.4$) because low-frequency cosine features approximate similar polynomials regardless of bandwidth. Within-scale correlations (diagonal) consistently exceed cross-scale correlations for all scale pairs. **(b)** Comparison of the naive diagonal variance estimate $\sum_k \mathrm{Var}(z_k)^2$ (blue) with the full within-scale Frobenius norm $\|\mathbf{\Sigma}_\ell\|_F^2$ (red) for each scale. Numbers indicate the correction ratio. At the finest scale ($\sigma_1$) the correction is minimal ($1.1\times$), but at coarse scales ($\sigma_5$–$\sigma_7$) it reaches 8–15$\times$, demonstrating that within-scale feature correlations must be accounted for in the Satterthwaite null distribution (Supplementary Note 3).

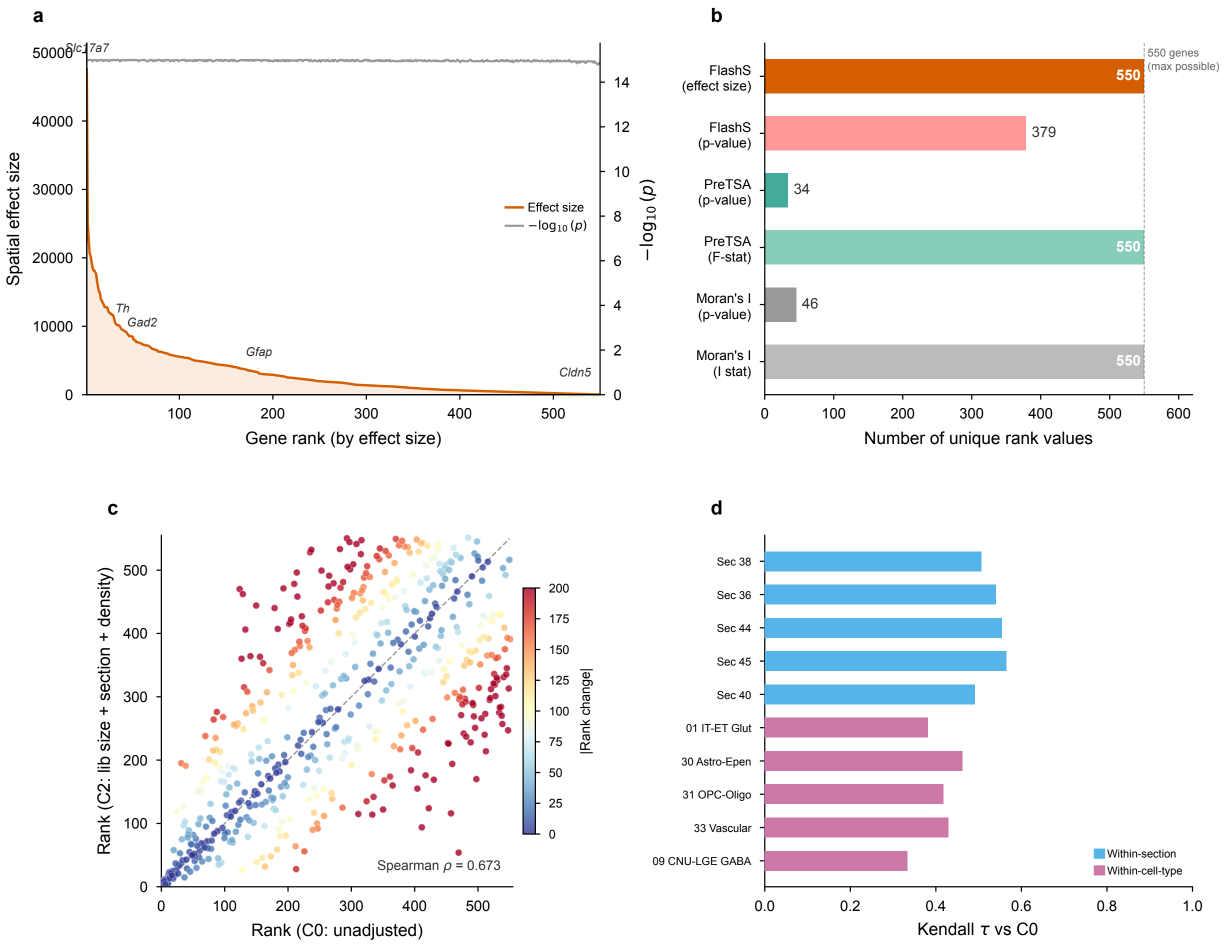

Figure 6: **Supplementary Figure 6: MERFISH atlas validation—ranking resolution and confounding robustness. (a)** Effect size versus p-value saturation for all 550 genes in the Allen MERFISH atlas (3.94 million cells), sorted by spatial effect size. The spatial effect size (orange) spans a broad dynamic range across the ranking, whereas $-\log_{10}(p)$ (grey) quickly saturates near machine precision for strongly spatial genes. Selected cell-type markers are annotated on the effect-size curve. **(b)** Number of unique rank values per method and ranking metric. FlashS's spatial effect size and PreTSA's F-statistic both achieve 550 unique values; p-value-only rankings collapse severely (PreTSA: 23; Moran's I: 46). **(c)** Rank stability across covariate adjustment. Scatter plot of gene ranks under the unadjusted model (C0) versus the fully adjusted model (C2: log library size + section fixed effects + local cell density) for all 550 genes. Points near the diagonal indicate stable rankings; color encodes the absolute rank change. Spearman $\rho = 0.67$ confirms moderate-to-strong concordance, with the largest rank shifts concentrated among mid-ranked genes. **(d)** Stratified Kendall $\tau$ computed by running FlashS independently within individual brain sections (blue) and cell types (pink), each compared against the full-dataset C0 ranking. Positive concordance within all strata confirms that spatial patterns persist within homogeneous subpopulations.

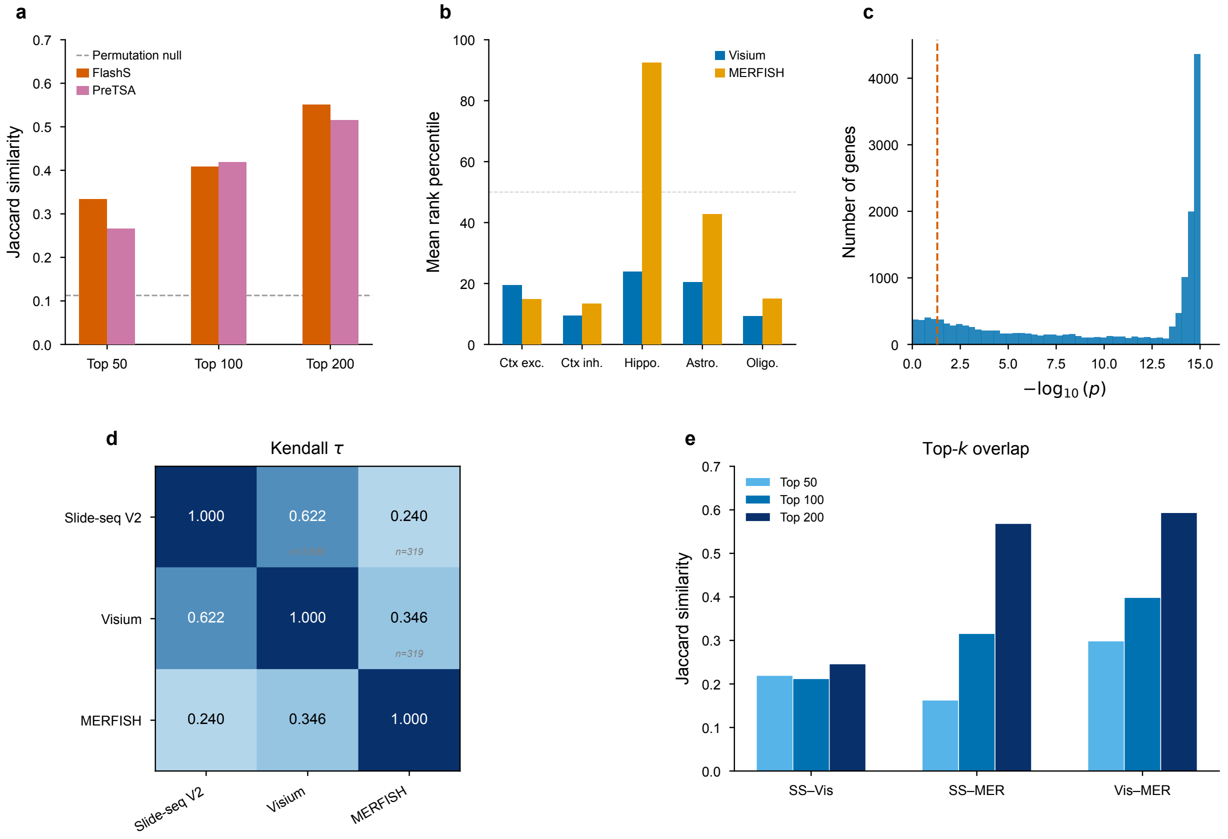

Figure 7: **Supplementary Figure 7: Cross-platform SVG ranking reproducibility. Row 1—Pairwise Visium–MERFISH validation: (a)** Top-$k$ Jaccard similarity between cross-platform SVG rankings for FlashS and PreTSA at $k = 50$, 100, and 200 (rank scatter shown in Fig. 4d), with the permutation null baseline (dashed line). **(b)** Brain region marker enrichment across platforms. Mean rank percentile of known markers for five cell populations on Visium and MERFISH data. **(c)** $-\log_{10}(p)$ histogram for 15,705 genes tested by FlashS on the 10x Visium V1_Adult_Mouse_Brain dataset (2,702 spots). Dashed red line marks $p = 0.05$. **Row 2—Three-platform extension (Visium, Slide-seq V2, MERFISH): (d)** Pairwise Kendall $\tau$ of FlashS SVG rankings across three platforms. Visium and Slide-seq V2 show strong concordance ($\tau = 0.622$, 3,666 common genes). Cross-modality comparisons show moderate concordance (Visium–MERFISH $\tau = 0.346$; Slide-seq–MERFISH $\tau = 0.240$; 319 common genes). **(e)** Top-$k$ Jaccard similarity across platform pairs at $k = 50$, 100, and 200. The brain marker gene rank heatmap is shown in Fig. 4f (main text).

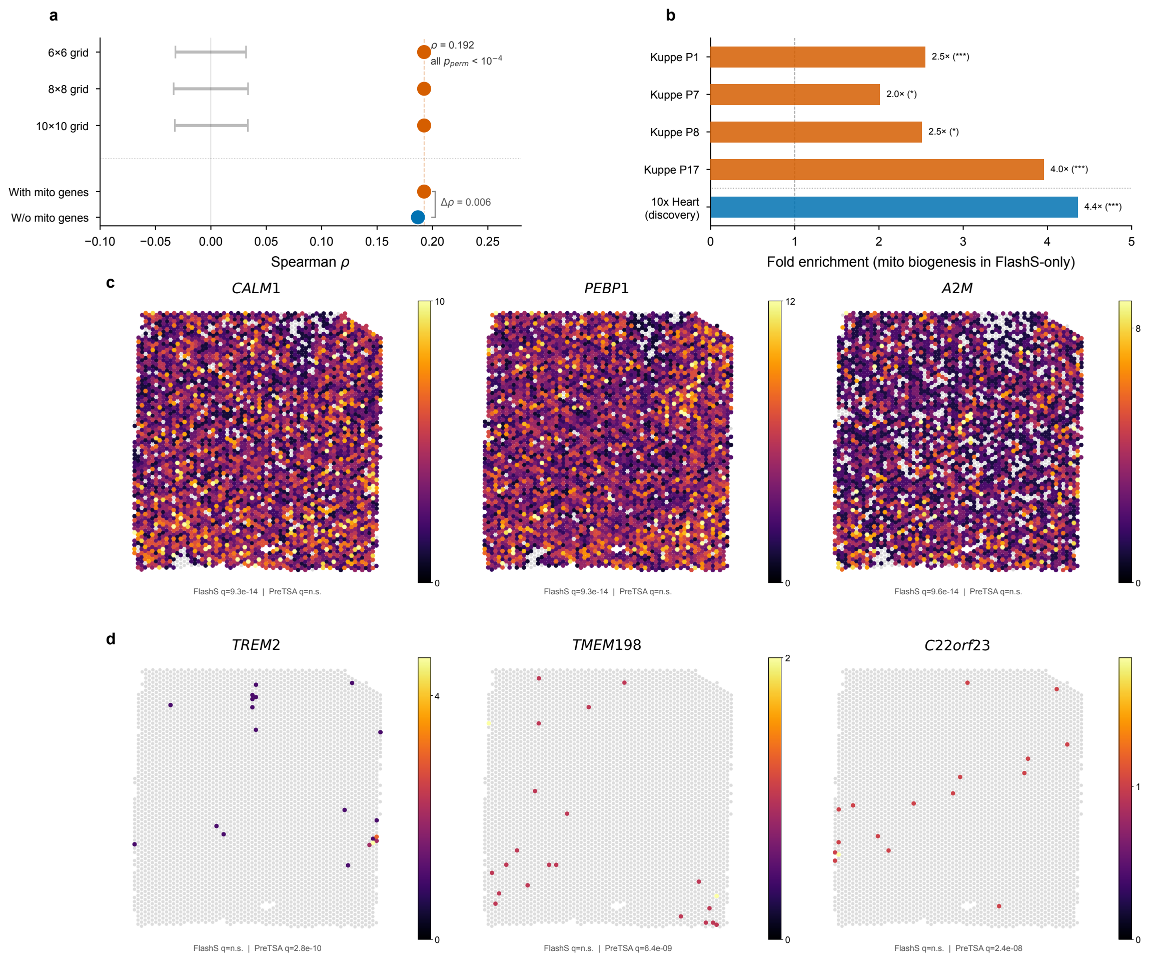

Figure 8: **Supplementary Figure 8: Heart case study validation.** **(a)** Block permutation and gene-set leakage controls for the mito–cardiomyocyte co-localization (main text Fig. 5g). Observed Spearman $\rho$ (vermillion dots) between the mitochondrial SVG module score and vCM proportion far exceeds the block-permutation null 95% CI (gray bands) across three spatial grid resolutions ($6 \times 6$, $8 \times 8$, $10 \times 10$; 10,000 permutations each; $p_{\text{perm}} < 10^{-4}$). Leakage-free control: diamond shows the original correlation; circle shows the correlation after removing all 49 mito biogenesis genes from the expression matrix prior to deconvolution ($\Delta\rho = 0.006$), confirming the co-localization is not driven by shared gene features. **(b)** Mitochondrial biogenesis fold enrichment in FlashS-unique SVGs across four healthy control Visium slides from Kuppe et al. (2022) and the original 10x Heart discovery dataset. Despite only 16 of 49 mito biogenesis genes being present in the Kuppe panel, enrichment is significant in all 4 replication samples (fold = 2.0–4.0, $p < 0.02$). Stars: hypergeometric $p$ (***$p < 0.001$). The cross-cohort correlation forest plot is shown in Fig. 5i. **(c)** Top 3 FlashS-unique SVGs (significant by FlashS at $q < 0.05$ but not by PreTSA), ranked by spatial effect size. All three genes (*CALM1*, *PEBP1*, *A2M*) show clear spatial expression patterns across the cardiac tissue. *CALM1* (calmodulin 1) is a calcium-signaling gene with known regional expression in cardiomyocytes; *PEBP1* encodes a Raf kinase inhibitory protein implicated in cardiac signal transduction. **(d)** Top 3 PreTSA-unique SVGs (significant by PreTSA at $q < 0.05$ but not by FlashS). These genes (*TREM2*, *TMEM198*, *C22orf23*) show sparse expression limited to isolated foci, consistent with cell-type-specific rather than tissue-wide spatial patterns. Adjusted p-values are shown below each panel.

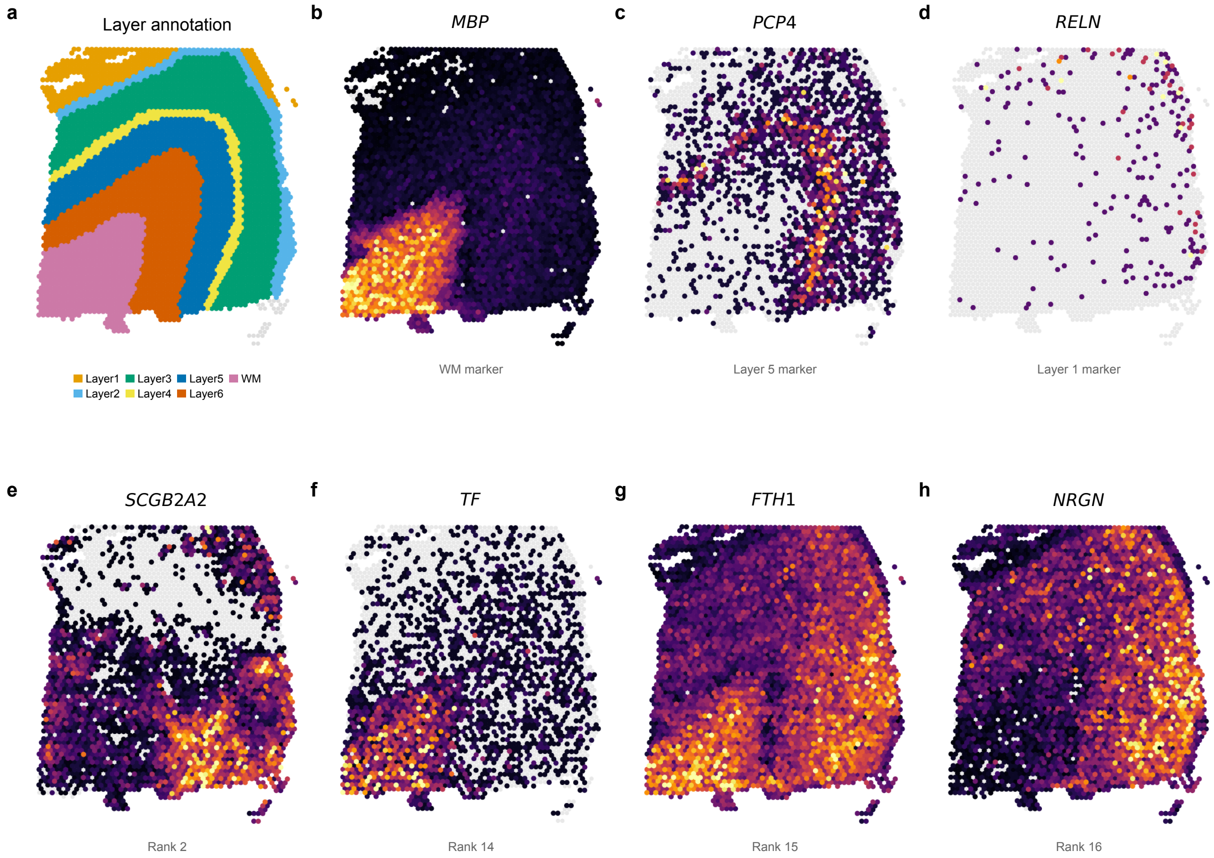

Figure 9: **Supplementary Figure 9: Spatial expression of** FlashS **SVGs on human DLPFC with cortical layer annotations. (a)** Manual cortical layer annotations for DLPFC Visium sample 151673 (Maynard et al. 2021; 3,639 spots, 7 layers). **(b–d)** Known cortical layer markers: *MBP* (white matter), *PCP4* (layer 5), and *RELN* (layer 1). Expression patterns match the annotated layer boundaries in (a). **(e–h)** Top-ranked non-classical FlashS SVGs: *SCGB2A2* (rank 2, localized expression), *TF* (transferrin, rank 14, WM-enriched gradient), *FTH1* (ferritin heavy chain, rank 15, cortical depth gradient), and *NRGN* (neurogranin, rank 16, gray-matter enriched). These genes are not among the 27 curated layer markers from Maynard et al. yet show clear layer-associated spatial patterns, demonstrating that FlashS identifies biologically relevant spatial variation beyond known markers.

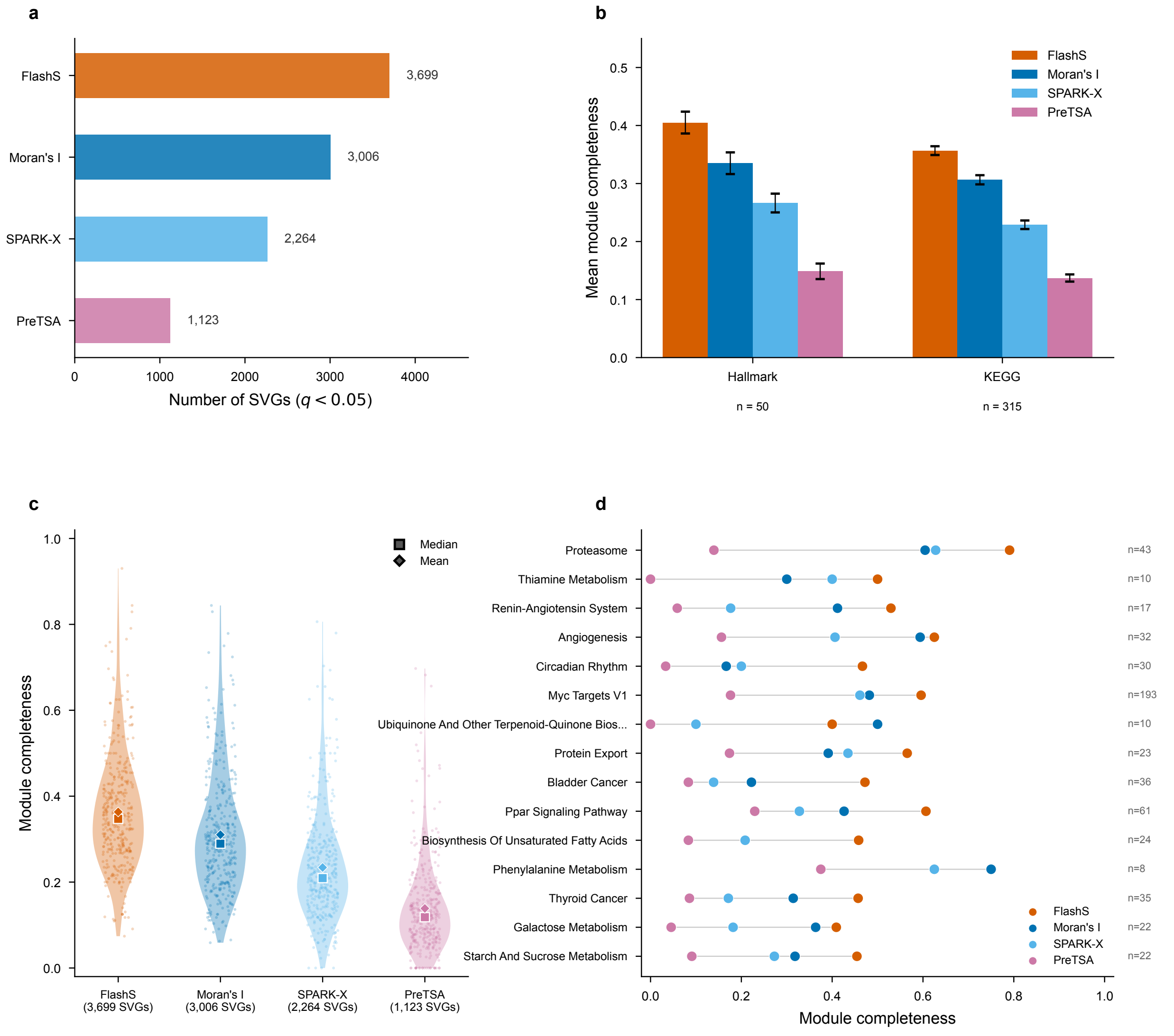

Figure 10: **Supplementary Figure 10: Extended module completeness analysis on Visium human heart.** Summary statistics are shown in Fig. 5f (main text) and Supplementary Table 12. All four methods are compared within the same 14,634-gene universe. **(a)** Number of SVGs detected by each method at $q < 0.05$ within the 14,634-gene universe, providing context for interpreting module completeness. **(b)** Mean module completeness (fraction of pathway genes detected as SVGs) for four methods across 50 MSigDB Hallmark and 315 KEGG pathways. Error bars indicate standard error of the mean. FLASHS achieves the highest pathway coverage (mean 0.363), followed by Moran's I (0.310), SPARK-X (0.234), and PreTSA (0.139). **(c)** Distribution of per-pathway module completeness across all 365 gene sets. Each dot represents one pathway; squares mark medians; diamonds mark means. SVG counts are shown below each method name. **(d)** Top 15 pathways with the largest FLASHS–PreTSA completeness gap. Each dot indicates the module completeness for one method; gray lines connect the range across methods. Pathway gene counts ($n$) annotated at right. The proteasome pathway shows the largest gap (FLASHS 0.79 vs PreTSA 0.14), reflecting its spatially organized expression that requires multi-scale kernel detection.

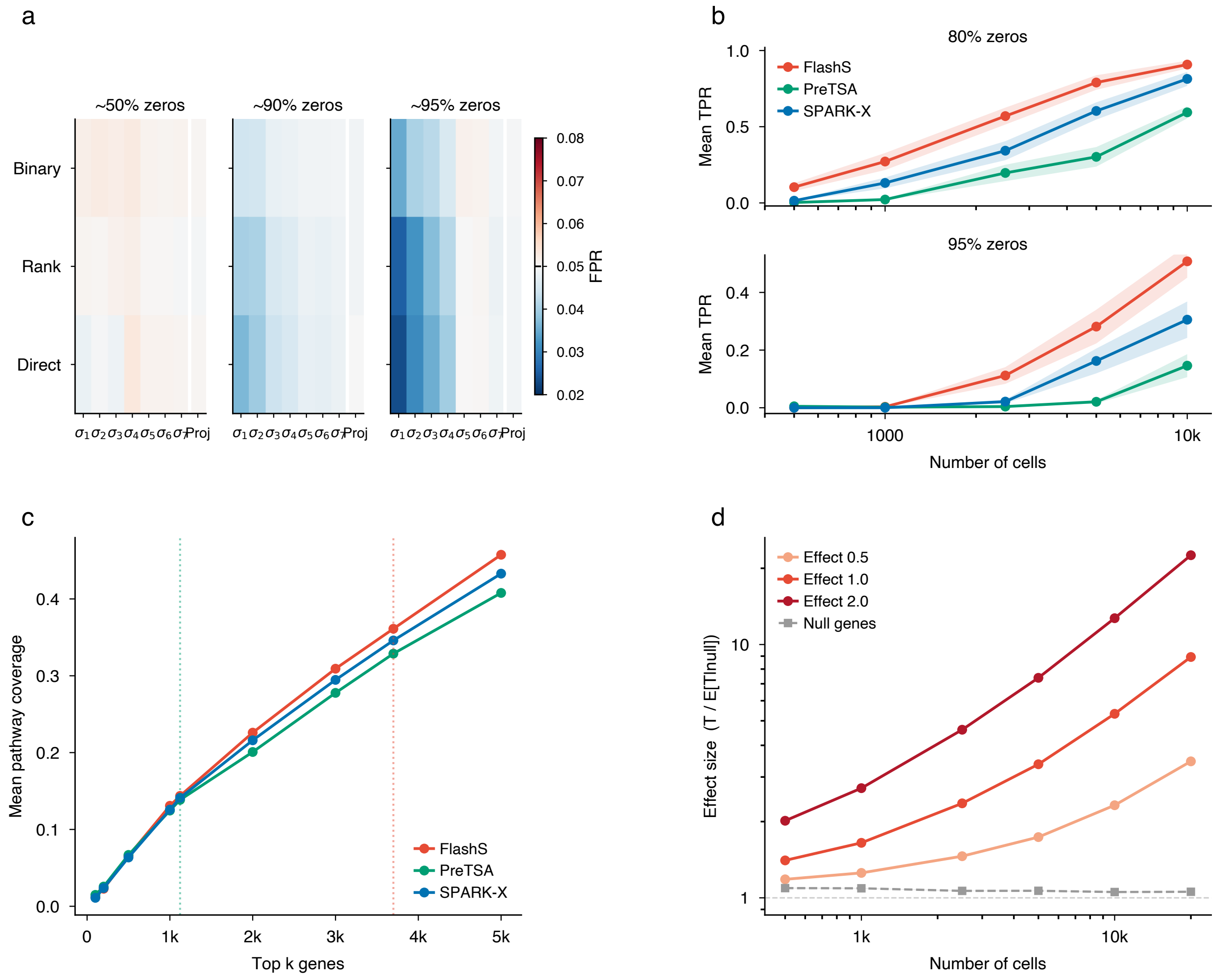

Figure 11: **Supplementary Figure 11: Extended statistical characterization of FlashS.** **(a)** Per-scale per-channel false-positive rate under the null (no spatial signal). Each cell shows the empirical FPR at $\alpha = 0.05$ averaged over 20 replicates (2,500 spots $\times$ 1,000 genes). The 24 individual kernel channels (3 encoding types $\times$ 7 bandwidth scales + 3 linear trend tests) all maintain FPR near the nominal level across sparsity regimes (50%, 90%, 95% zeros), confirming that the Cauchy combination does not introduce inflation at any component level. **(b)** Statistical power (true-positive rate at $q < 0.05$, BH-corrected) versus sample size for FlashS, PreTSA, and SPARK-X. Results are averaged across four spatial patterns (gradient, hotspot, multiscale, domain) with effect size 1.0 and 10 replicates per condition. FlashS achieves consistently higher power, with the gap widening at high sparsity (95% zeros). **(c)** Top-$k$ controlled module completeness on Visium human heart. For each method, the top $k$ genes by score are selected and mean pathway coverage (fraction of MSigDB Hallmark + KEGG gene sets recovered) is computed. Vertical dashed lines indicate the number of SVGs detected by PreTSA ($k = 1{,}123$) and FlashS ($k = 3{,}699$) at $q < 0.05$. **(d)** Effect-size convergence with increasing sample size. The multi-kernel test statistic ratio $T/\mathbb{E}[T \mid \text{null}]$ grows monotonically for signal genes (three effect sizes shown) while remaining near 1.0 for null genes, demonstrating consistent statistical power with well-calibrated null behavior (80% sparsity, gradient pattern, 10 replicates).

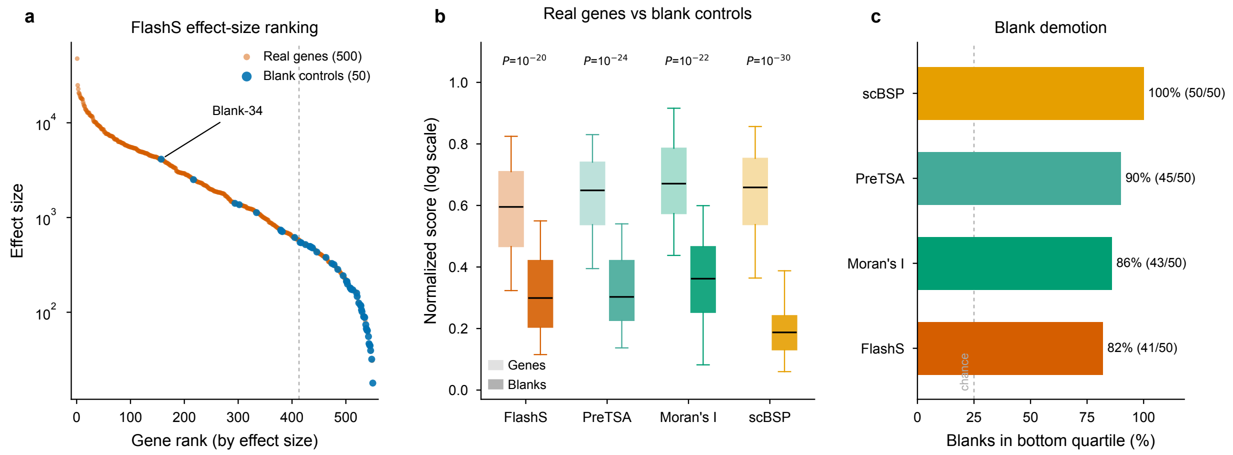

Figure 12: **Supplementary Figure 12: MERFISH blank negative-control ranking across SVG methods.** **(a)** FlashS effect-size ranking of all 550 genes in the Allen MERFISH atlas (3.94 million cells). Real genes (500, vermillion) and blank negative-control barcodes (50, blue) are shown. Blanks cluster at the bottom of the ranking (median rank 505.5/550, 96% in the bottom half), indicating that the spatial effect size separates assayed genes from blank controls even when p-values saturate at this sample size. One outlier (Blank-34, rank 157) may reflect spatially structured technical signal. Dashed line marks the bottom-quartile cutoff (rank 413). **(b)** Distribution of normalized spatial scores for real genes (light boxes) versus blank controls (dark boxes) across four SVG detection methods. All four methods assign significantly lower scores to blanks (Mann–Whitney $U$ test $p$-values annotated). Scores are min-max normalized on the $\log_{10}$ scale for visual comparability across methods with different score ranges. **(c)** Percentage of blank controls ranked in the bottom quartile by each method. All methods exceed the 25% chance level (dashed line), with scBSP achieving 100% blank demotion. FlashS places 82% of blanks in the bottom quartile.
